# Hybridization dynamics and extensive introgression in the *Daphnia longispina* species complex: new insights from a high-quality *Daphnia galeata* reference genome

**DOI:** 10.1101/2021.02.01.429177

**Authors:** Jana Nickel, Tilman Schell, Tania Holtzem, Anne Thielsch, Stuart R. Dennis, Birgit C. Schlick-Steiner, Florian M. Steiner, Markus Möst, Markus Pfenninger, Klaus Schwenk, Mathilde Cordellier

## Abstract

Hybridization and introgression are recognized as an important source of variation that influence adaptive processes; both phenomena are frequent in the genus *Daphnia,* a keystone zooplankton taxon in freshwater ecosystems that comprises several species complexes. To investigate genome-wide consequences of introgression between species, we provide here the first high-quality genome assembly for a member of the *Daphnia longispina* species complex, *Daphnia galeata*. We further re-sequenced 49 whole genomes of three species of the complex and their interspecific hybrids both from genotypes sampled in the water column and from single resting eggs extracted from sediment cores. Populations from habitats with diverse ecological conditions offered an opportunity to study the dynamics of hybridization linked to ecological changes and revealed a high prevalence of hybrids. Using phylogenetic and population genomic approaches, we provide first insights into the intra- and interspecific genome-wide variability in this species complex and identify regions of high divergence. Finally, we assess the length of ancestry tracts in hybrids to characterize introgression patterns across the genome. Our analyses uncover a complex history of hybridization and introgression reflecting multiple generations of hybridization and backcrossing in the *Daphnia longispina* species complex. Overall, this study and the new resources presented here pave the way for a better understanding of ancient and contemporary gene flow in the species complex and facilitate future studies on resting egg banks accumulating in lake sediment.

## Introduction

Gene flow between species can be pervasive and can affect substantial parts of the genome. Hybridization and introgression are recognized as an important source of variation that can influence adaptive processes in plants, animals, yeast, and fungi (reviewed in Abbott*, et al.* 2013; Arnold and Martin 2009). The amount of realized gene flow varies among taxa and along the genome; it is governed by intrinsic genomic features such as recombination rate, structural variation and intrinsic incompatibilities, as well as the species’ biology and ecology including ecological and sexual selection, migration, and mode of reproduction.

How can species in diversifying clades frequently hybridize and show introgression but nevertheless maintain species boundaries? A growing body of literature provides examples for a high variety of systems where speciation occurs in the face of gene flow (e.g. Fraïsse*, et al.* 2014; Martin*, et al.* 2019; Meier*, et al.* 2017). However, it is important to recognize that these systems are distributed along a wide spectrum. On one side of this spectrum, hybridization occurs but is not followed by introgression for several reasons such as reduced hybrid fertility or strong selection against hybrid phenotypes, leading to rapid hybrid breakdown. Barth*, et al.* (2020) found that species boundaries in tropical eels are stable despite millions of years of hybridization, and also observed very few admixed individuals beyond F1 and first-generation backcrosses. The hybrid breakdown observed in this system reduces the likelihood of introgression via backcrossing. On the other side of the spectrum, hybridization is followed by introgression, and ongoing exchange of genetic information between species (e.g. Butlin*, et al.* 2014; Doellman*, et al.* 2018; Kaiser*, et al.* 2021; Martin*, et al.* 2013). Several empirical studies (Canestrelli*, et al.* 2017; Schreiber and Pfenninger) as well as theoretical models (Flaxman*, et al.* 2014; Rafajlović*, et al.* 2016; Yeaman and Whitlock 2011) suggest the possibility of intermediate constant equilibrium states, meaning that certain parts of the genome remain diverged (‘islands’ or ‘continents of divergence’), while others are freely exchanged among closely related species without ever reaching complete genomic isolation.

Recurrent hybridization and introgression are frequent in the genus *Daphnia* (Crustacea, Cladocera) Members of the genus have served as ecological model organisms for over a century (e.g. Miner*, et al.* 2012), and the first crustacean genome to be sequenced was that of a member of the *Daphnia pulex* species complex (Colbourne*, et al.* 2011). Since then, the genomes of 45 crustaceans have been sequenced with a focus on species of economic or medical interest (NCBI, last accessed January 2021). Despite their key role in marine and freshwater food webs around the globe, genomic resources for zooplanktonic species are still scarce. In many aquatic food webs, zooplanktonic crustaceans link primary production by phytoplankton and secondary consumers, such as planktivorous fish and larger invertebrate species (Lampert and Sommer 2007). (e.g. Gannon and Stemberger 1978; Gliwicz 1990)

*Daphnia* are highly phenotypically plastic and a textbook example for inducible defense mechanisms (Tollrian and Harvell 1999), as they respond to variation in predation risk through spectacular changes in morphology. Further, *Daphnia* are cyclical parthenogens and hence able to alternate between asexual and sexual reproduction. They reproduce asexually through longer periods of time, and the product of sexual reproduction events (usually seasonal) are resting eggs able to withstand adverse conditions for decades and even centuries (Frisch*, et al.* 2014). Resting eggs extracted from sediment cores can be hatched, and ancient genotypes brought to life (reviewed in Orsini*, et al.* 2013). Moreover, the DNA preserved in those resting eggs can be directly analyzed with various molecular methods (e.g. Cousyn*, et al.* 2001; Dziuba*, et al.* 2020; Lack*, et al.* 2018). Thus, cyclical parthenogenesis, biological archives in lake sediments and high levels of phenotypic plasticity make *Daphnia* a particularly interesting model for evolutionary studies.

The genus *Daphnia* is composed of two subgenera, *Ctenodaphnia* and *Daphnia*, and two groups are delimited within the subgenus *Daphnia*: the *D. pulex* group *sensu lato* and the *D. longispina* group *sensu lato* (see Adamowicz*, et al.* 2009). The latter is sometimes also referred to as subgenus *Hyalodaphnia* and includes the *Daphnia longispina* species complex (DLSC) (Petrusek*, et al.* 2008a). The two *Daphnia* groups are highly differentiated and share their most recent common ancestor around 30 Mya (MRCA *D. longispina* – *D. pulex* group, MRCA *D. longispina* – *D. pulex* group, Cornetti*, et al.* 2019). Members of the genus *Daphnia* show little variation in chromosome number, with most species having 10 pairs of chromosomes, except for the *D. pulex* group with n=12 (Beaton and Hebert 1994; Trentini 1980). All sequenced and assembled *Daphnia* genomes so far belong either to the *D. pulex* group or the subgenus *Ctenodaphnia*, however no high-quality reference genome of the third major group, the *D. longispina* group (*Hyalodaphnia*) is published.

The prevalence of hybridization in the genus *Daphnia* across taxa and ecosystems and its impact on their evolutionary history has intrigued researchers for decades (e.g. Schwenk 1993; Vergilino*, et al.* 2011; Wolf 1987). In contrast to many other well-studied hybrid systems (Barton and Hewitt 1985) with clear defined hybrid zones where species’ ranges overlap, the distribution of *Daphnia* species and their hybrids is more of a fragmented nature: they occupy lake and pond ecosystems that vary in their ecological characteristics and hence constitute a mosaic across the landscape. Ecologically differentiated taxa and their hybrids are thus distributed across habitat patches (Harrison 1986). Within these patches, the possibility to interrogate biological archives also revealed fluctuations in *Daphnia* community composition over time (e.g. Alric*, et al.* 2016; Brede*, et al.* 2009), associated with hybridization events among species in some cases. Variation in hybridization events across time and among habitats has often been observed in correlation with ecological changes, such as eutrophication or global change (Brede*, et al.* 2009; Cordellier*, et al.* 2021; Dziuba*, et al.* 2020; Keller*, et al.* 2008; Rellstab*, et al.* 2011; Spaak*, et al.* 2012).

Members of the *Daphnia longispina* species complex inhabit many large ponds and lakes in central and northern Europe, and three of them have been particularly well studied: *Daphnia galeata*, *Daphnia longispina* and *Daphnia cucullata* (Petrusek*, et al.* 2008a). These species can coexist, but earlier studies suggest gene flow among them is limited (Spaak 2004). Despite their obviously ancient divergence (Schwenk*, et al.* 2000), DLSC species are still able to form interspecific hybrids, although not all combinations are equally likely to lead to viable and fertile individuals (Schwenk*, et al.* 2001). A mechanism preventing gene flow among species might be their different ecological preferences, e.g., regarding trophic level (Spaak*, et al.* 2012), food quality (Seidendorf*, et al.* 2007), and predation pressure (Spaak and Hoekstra 1997);(Petrusek*, et al.* 2008b).

Up to now, genetic markers available to study hybridization in the DLSC are limited to allozymes (Wolf and Mort 1986), a few mitochondrial regions (Schwenk 1993), a dozen microsatellite markers (Brede*, et al.* 2006; Thielsch*, et al.* 2012) and a few further nuclear loci (Billiones*, et al.* 2004; Rusek*, et al.* 2015; Skage*, et al.* 2007). Seminal studies such as Brede*, et al.* (2009) and Limburg and Weider (2002) first made use of microsatellite markers to analyze environmentally driven shifts in allelic frequencies, species and hybrid composition of the DLSC communities in Lake Constance and Belauer See over time, respectively. Further, a number of studies addressed the spatial distribution of DLSC species/taxa with these markers (e.g. Griebel*, et al.* 2016; Ma*, et al.* 2019; Thielsch*, et al.* 2017). These low-resolution markers allowed to identify hybrid individuals and brought evidence for introgression but could not provide the resolution necessary to either assess how pervasive introgression is or how it varies across the genome. Further, it is not clear whether introgression occurs among all three species to the same extent. Given the ubiquitous hybridization among the DLSC taxa, the question also arises why they are still well distinguishable species. Whether the DLSC represents a case of incipient speciation, introgression after secondary contact, speciation reversal, or has reached an intermediate constant equilibrium state, among other possibilities, can only be answered with genome-wide analyses empowered by a high-quality genome assembly.

Here, we present a high-quality assembly for *Daphnia galeata*, thus filling an important gap for *Daphnia* whole-genome studies. Furthermore, to facilitate genome-wide assessments of divergence across species and of introgression between species, we conducted genome-wide resequencing studies in the DLSC. We analyzed whole-genome sequences of parental species and their interspecific hybrids, both from genotypes obtained in the wild and maintained in laboratories, and from single resting eggs extracted from sediment cores. We provide first insights into the intra- and interspecific genome-wide variability in this species complex and identify regions of high divergence. We reconstructed the phylogenetic relationships in the species complex using whole mitochondrial genomes. Finally, we assess the length of ancestry tracts in different classes of hybrids to characterize introgression patterns. Our study paves the way for long-awaited analyses on the dynamics of introgression in this complex and exploitation of the unique opportunity this group has to offer: a window of more than one hundred years of evolution in action.

## Results

### Genome assembly

The raw assembly was obtained by combining PacBio long reads (1,679,290, 11.52 Gb) and Illumina short reads (70,310,338, 9.79 Gb after trimming) and using the hybrid assembler RA (https://github.com/lbcb-sci/ra). It originally comprised 1,415 contig sequences covering a total length of 153.6 Megabases (Mb), with an N50 value of 172 kilobases (kb) and a slightly elevated GC content (40.02% Supplementary Methods Table 3) compared with the values expected for a *Daphnia* species (see Table 1). According to an analysis based on coverage and GC content of the contig sequences conducted with blobtools (Laetsch and Blaxter 2017), a portion of the assembly consisted of non-*Daphnia* contigs, which could then be removed (267 contigs, equaling 22.97 Mb). Consequently, GC content decreased to 38.75%, nearing the values obtained for other *Daphnia* assemblies (see Table 1 for an overview). The application of this filter as well as the exclusion of the mitochondrial genome led to a decrease in the number of sequences and the total length of the assembly. Iterative scaffolding led to a decrease in the total number of sequences. This together with a substantial increase in N50 resulted in a highly contiguous assembly, with a total length of 133,304,63 basepairs (bp), an N50 of 756.7 kb and only 346 sequences, i.e., on average 30 sequences per chromosome. Contiguity statistics for the different assembling steps are given in Supplementary Methods Table 3.

**Table 1:** Assembly metrics and annotation statistics for the present assembly and two previously published *Daphnia* assemblies. Contiguity statistics of the annotation were calculated excluding tRNAscan results. BUSCO 3.0.2 was executed in protein mode for the different MAKER rounds. Conserved Domain Arrangements (CDAs) were searched with Pfam scan 1.6 and DOGMA 3.4. Results for BUSCO and DOGMA completeness statistics are given in percent.

| Species | *D. galeata* | *D. pulex* (Ye *et al*) | *D. magna* (Lee *et al*) |
| --- | --- | --- | --- |
| Strain | M5 | PA42 | SK |
| **Assembly metrics** |  |  |  |
| # scaffolds | 346 | 493 | 4,192 |
| Largest scaffold (bp) | 2,950,711 | 7,584,612 | 16,359,456 |
| Total length (bp) | 133,304,630 | 189,550,516 | 122,937,721 |
| N50 (bp) | 756,671 | 1,160,003 | 10,124,675 |
| L50 (bp) | 48 | 36 | 5 |
| GC (%) | 38.75 | 40.39 | 40.54 |
| # N's | 120,845 | 4,006,006 | 82,97,703 |
| # N's per 100 kbp | 90.65 | 2113.42 | 6749.52 |
| **Annotation** |  |  |  |
| **Number** |  |  |  |
| Gene | 15,845 | 18,440 | 15,721 |
| mRNA | 16,774 | 18,440 | 15,721 |
| Exon | 117,364 | 128,688 | 95,203 |
| CDS | 119,402 | 118,916 | 94,047 |
| **Mean** |  |  |  |
| mRNAs/gene | 1.06 | 1 | 1 |
| Exons/mRNA | 7.00 | 6.98 | 6.06 |
| CDSs/mRNA | 7.12 | 6.45 | 5.98 |
| **Median length (bp)** |  |  |  |
| Gene | 2,097 | 1,919.5 | 1,586 |
| mRNA | 2,142 | 1,919.5 | 1,521 |
| Exon | 167 | 162 | 160 |
| Intron | 74 |  |  |
| CDS | 152 | 144 | 159 |
| **Total space (bp)** |  |  |  |
| Gene | 51,689,473 | 53,936,938 | 37,505,261 |
| mRNA | 51,689,329 | 53,936,938 | 36,178,687 |
| Exon | 29,314,592 | 30,208,483 | 22,336,755 |
| CDS | 25,132,876 | 23,586,918 | 21,881,778 |
| **Single** |  |  |  |
| Exon mRNA | 663 | 144 | 1,775 |
| CDS mRNA | 710 | 0 | 0 |
| **BUSCO N=1066** |  |  |  |
| C | 94.3 | 94.1 | 97.0 |
| S | 91.7 | 82.6 | 95.3 |
| D | 2.6 | 11.5 | 1.7 |
| F | 0.7 | 3.5 | 1.7 |
| M | 5.0 | 2.4 | 1.3 |
| DOGMA N=4222 | 93.63 | 91.43 | 93.91 |

Mapping the filtered Illumina reads with bwa mem (Li 2013) and PacBio reads with Minimap 2.17 (Li 2018), resulted in a mapping rate of respectively 94.1% and 85.5%. The coverage distribution can be seen in Figure S1B.

According to blobtools results, no contamination could be identified in the final assembly (Figure S1A). Remaining scaffolds (12, amounting to a total length 1.79Mb) with taxonomic assignment other than Arthropoda were kept because coverage and GC are similar to *D. galeata* scaffolds and taxonomic assignment alone might be false positive. Further, the completeness assessment through BUSCO (Simão*, et al.* 2015, Arthropoda set, odb9) indicated 95.7% of complete single copy core orthologs and a very low duplication rate (C: 95.7% [S: 94.7%, D: 1.0%], F: 0.8%, M: 3.5%, n: 1066). The genome size was estimated based on mapped nucleotides and mode of the coverage distribution by backmap 0.3 (https://github.com/schellt/backmap), resulting in 156.86Mb and 178.03Mb for Illumina (52x) and PacBio (26x) respectively, and by k-mer based approach using GenomeScope resulting in a size of 150.6Mb.

When compared to other published full genomes for *Daphnia* species, the *D. galeata* final assembly is shorter than both *D. pulex* assemblies (Colbourne*, et al.* 2011; Ye*, et al.* 2017), and roughly the same size as *D. magna* (Lee*, et al.* 2019), which also has 10 chromosomes (Table 1). The GC content is lower, which can be attributed to the strict filtering for contamination applied pre- and post-assembly, a procedure not applied in the other *Daphnia* assemblies, to our knowledge. Even though Lee *et al.* (2019) and Ye *et al.* (2017) treated the animals with antibotics before sequencing this suggests that these genome assemblies contain more contigs of bacterial origin than the *D. galeata* assembly. Thanks to the use of long-read data, iterative scaffolding and gap filling, the number and length of assembly gaps (Ns) is substantially lower and contiguity is high (but see Table 3 in Supplementary Methods).

### Genome annotation

After applying RepeatMasker (Smit*, et al.* 2013-2015) with the custom repeat library described in the methods section, 21.9% of the assembly was masked. The distribution of masked fraction per repeat element can be found in Supplementary Methods Table 5.

The final annotation with MAKER (Holt and Yandell 2011) predicts 15,845 genes with a median length of 2,097 base pairs. There is an average of 1.06 mRNAs per gene and 7 exons/mRNA (Table 1). The total number of predicted mRNA substantially differs from the number of transcripts previously published for this species (32,903, Huylmans*, et al.* 2016). This is not surprising, as this transcriptome assembly did not make use of protein evidence we included here, and might contain isoforms. Further, it was based on a pool of mRNA from different clonal lines, and the assembly process might have been impeded by allelic diversity. As further quality criterion, the Annotation Editing Distance (AED) was compared across the three MAKER rounds and is visualized in Supplementary Methods Figure 4. AED improved mostly between rounds 1 and 2 of the annotation but only marginally with a further round.

A high percentage of protein sequences could be annotated: 15,898 (94.78%) with InterProScan (Jones*, et al.* 2014) and 15,960 (95.15%) with blast against Swiss-Prot. With this combination of searches, a hit within InterProScan and blast was found for 16,675 protein sequences (99.41%). GeneOntology annotation was possible for 9555 sequences (56.96%). A detailed overview of the functional annotated sequences per database or search algorithm is shown in Supplementary Methods Table 7.

### Genotyping

Short-read sequence data were generated for 72 individuals: 17 unamplified DNA samples from isofemale clonal lines and 55 whole genome amplification (WGA) samples (conducted on single resting eggs) that passed PCR contamination checks. After screening for contamination and removing datasets with only very few reads mapping to the *D. galeata* reference, 49 single genotypes remained: 32 from resting eggs and 17 from clonally propagated lines, established from individuals sampled in the water column or hatched from resting eggs (Figure 1A, Table S2). Data gained from clonal lines with a species attribution were used as “parental species” data: five samples for *D. galeata*, four for *D. longispina*, and three for *D. cucullata*. The parental clones are part of two larger clone panels representing the parental species and their diversity in several European lakes. Their identity was established prior to this study either based on mitochondrial and microsatellite markers (M5, LC3_6, J2, Herrmann*, et al.* 2017) or morphological examination, mitochondrial markers and factorial correspondence analyses based on microsatellite markers (Alric*, et al.* 2016; Möst 2013), including hybrids and historical resting eggs, which separates parental species and hybrids (e.g. Alric*, et al.* 2016; Dlouha*, et al.* 2010; Rellstab*, et al.* 2011; Yin*, et al.* 2014). In addition, data were available for four resting eggs from Arendsee (AR), 12 resting eggs from Dobersdorfer See (DOB), five clonal lines and eight resting eggs from Eichbaumsee (EIC), and eight resting eggs from Selenter See (SE) (Table S2). While the analysis of eggs from older sediment layers was attempted, biological material was either limited, of poor quality, or contaminated. Our isotope dating for DOB was unconclusive: either slides of the cored location or a high sedimentation rate meant the top 30 cm of the core didn’t show the usual isotope peaks, thus preventing precise dating. EIC samples were recent since they were collected from surface bank sand. For SE, the oldest eggs analyzed here originated from the 2-3cm layer of the core, which corresponds to max. ~17 years (pers. comm Thorbjørn Andersen). For AR the oldest eggs for which results were obtained were isolated from the 4-5cm layer, corresponding to ~2005 (pers. comm. Miklos Balínt).

**Figure 1:**
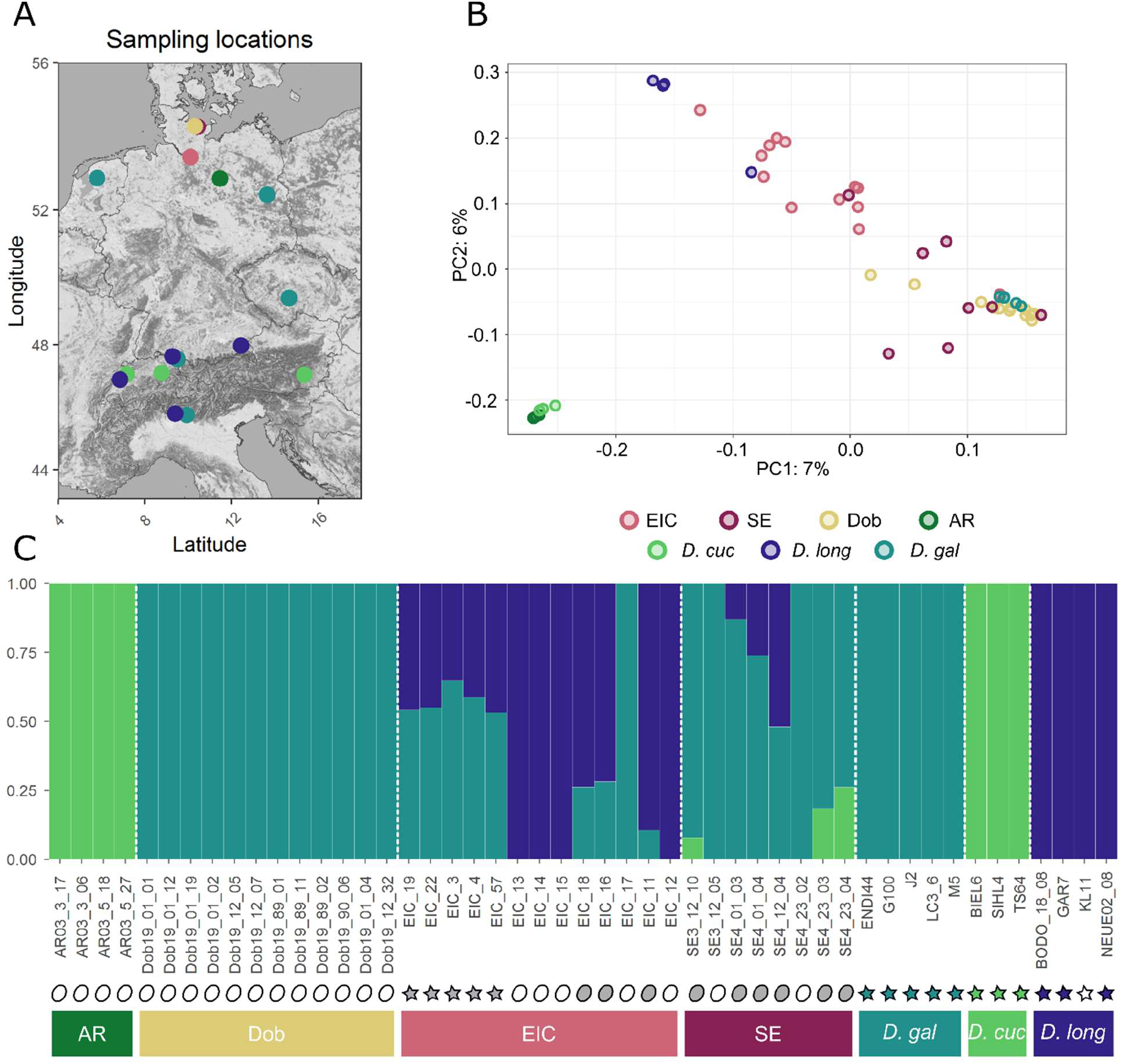
Parental species: *D. gal*: *D. galeata*, *D. long*: *D. longispina*, *D. cuc*; *D. cucullata*, populations: AR: Arendsee, Dob: Dobersdorfer See, SE: Selenter See, EIC: Eichbaumsee. Color coding is consistent throughout panels A and B **A.** Map of the sampling locations. **B.** PCA plot obtained with SNPrelate, including loci with linkage r^2^ <0.5 within 500-kb sliding windows. **C.** Admixture plot obtained with K=3. Symbols indicate the sample type: oval for genotypes sequenced directly from resting eggs, stars for genotypes sampled in the water column and propagated clonally in the laboratory prior to sequencing. Symbol filling indicates how these genotypes were classified in subsequent analyses: white for non-admixed genotypes, grey for admixed genotypes, green, blue and teal for genotypes used as representatives for parental species. Bottom bars are color coded to match the color scheme used in panels A and B.

An average of 89.9% (range: 31.7-98.6%) reads aligned to the reference genome with a mean coverage of 10.26x (range: 0.34-52.30x) (Table S3). The final SNP data set for subsequent analyses after quality-filtering included 3,240,339 SNPs across the 49 samples. To rule out possible reference bias we compared mapping rates of reads with the reference allele and to the alternative allele at heterozygous sites. We found no preferential mapping of the reference allele, as all species categories and the hybrids had a median distribution close to 0.5 (Figure S2).

### Principal Component Analysis

In a PCA including all genotypes and conducted with SNPRelate v1.20.1 (Zheng*, et al.* 2012), the parental species genotypes grouped in three very distinct clusters. *D. cucullata* separated from *D. galeata* along PC1, which explained 7% of the variation. *D. longispina* separated from *D. galeata* and *D. cucullata* along PC2 which explained 6% of the variation (Figure 1B). Although sampled in different lakes, all parental species genotypes were grouped in tight clusters along the two axes with little evidence for population substructure. Population AR clustered with the *D. cucullata* reference individuals while population samples from DOB, EIC and SE were more spread out, mostly between the *D. galeata* and *D. longispina* clusters.

### Admixture analyses uncover hybrids

The PCA results are confirmed by an admixture analysis conducted with ADMIXTURE (Alexander and Lange 2011) with K=3, supported by the lowest cross-validation error of the tested K values. The known parental species genotypes were clearly separated into three clusters (Figure 1C). While we detect no evidence of admixture in the AR and DOB samples and, based on our parental species, consider them to belong to the species *D. cucullata* and *D. galeata*, respectively, the two other populations seem to consist mostly of admixed individuals. The five EIC samples sequenced after clonal propagation were all found to be admixed (*D. galeata* and *D. longispina*), while the EIC resting eggs were either admixed (3) or belonged to one of the parental species (5). SE resting eggs present all combinations of admixture except *D. cucullata x D. longispina*: *D. galeata* (2), admixed between *D. galeata* and *D. cucullata* (3), and admixed between *D. galeata* and *D. longispina* (3).

### Ancestry painting

Based on results obtained in the ADMIXTURE analysis, two pairs of species and their putative hybrids were analyzed with an “ancestry painting” approach, outlined in Barth*, et al.* (2020) and Runemark*, et al.* (2018a): *D. galeata* and *D. longispina* parental genotypes and putative hybrids between them from populations EIC and SE, and *D. galeata* and *D. cucullata* parental genotypes and putative hybrids between them from population SE. Briefly, after identifying fixed sites for each of the species in the analyzed pair, heterozygosity was calculated for these sites and a hybrid index derived from the obtained results (https://github.com/mmatschiner/tutorials/tree/master/analysis_of_introgression_with_snp_data). Further, information on the maternal species is used to tentatively categorize the admixed individuals. For a first-generation hybrid (F1) the expectation would be 50% of the nuclear genome being derived from each parental species (hybrid index ≈ 0.5) and mostly heterozygous fixed sites (heterozygosity ≈ 1.0). Individuals originating from the backcrossing of F1 with one of the parental species are expected to have hybrid index values around 0.25 or 0.75. (Figure 2D). We consider individuals with intermediate hybrid indices (>0.25 and <0.75) and lower heterozygosity (<0.5) to be later-generation hybrids, meaning they have one or multiple hybrid ancestors we are not able to classify further (Slager*, et al.* 2020) We consider individuals with a hybrid index of ≤0.25 or ≥0.75 to be backcrossed with the respective parental species in at least one generation and the majority of the genome derives from one species. This definition is broad and will be refined with the addition of a greater number of parental genotypes.

The comparison of genotypes from parental species *D. galeata* and *D. longispina* (five and three individuals, respectively) allowed identifying a total of 335,052 fixed sites between the two species. Due to the quality filters applied to parental and hybrid genotypes, we could analyze 131,914 of these fixed sites in the putative hybrids, where read coverage was sufficient. The diploid genotypes were then plotted for all hybrids as homozygous for either of the parental species or as heterozygous (Figure 2A for the 50 longest scaffolds). The *D. longispina* reference clone KL11 was excluded from further analysis due to issues with missing data. All eleven genotypes from SE and EIC identified as likely *D. galeata* x *D. longispina* hybrids in the ADMIXTURE analysis possessed a *D. galeata* mitochondrial genome. The proportion of the maternal *D. galeata* genome in these hybrids, however, varied greatly, between 27.4% and 86.6%, and they all showed low heterozygosity, between 9.1% and 34.6% (Figure 2C, Table 2). These values are unlikely for F1 hybrids or backcrosses of F1 with one of the parental species (Table 2).

**Table 2:** Data derived from ancestry painting analysis and based on the fixed sites inferred from analyzing parental species genotypes. Maternal species attribution is based on mitochondrial phylogeny, hybrid attribution is based on the ADMIXTURE plot.

| Sample | Hybrid Index | Heterozygosity | Maternal species | Hybrid | Interpretation |
| --- | --- | --- | --- | --- | --- |
| SE_12_10 | 0.079 | 0.064 | cuc | gal x cuc | Backcross gal |
| SE_23_03 | 0.183 | 0.348 | cuc | gal x cuc | Backcross gal |
| SE_23_04 | 0.267 | 0.509 | cuc | gal x cuc | Unclear |
| EIC_19 | 0.653 | 0.156 | gal | gal x long | Later-generation |
| EIC_22 | 0.644 | 0.283 | gal | gal x long | Later-generation |
| EIC_3 | 0.751 | 0.095 | gal | gal x long | Backcross gal |
| EIC_4 | 0.693 | 0.144 | gal | gal x long | Later-generation |
| EIC_57 | 0.637 | 0.267 | gal | gal x long | Later-generation |
| EIC_18 | 0.393 | 0.302 | gal | gal x long | Later-generation |
| EIC_16 | 0.403 | 0.349 | gal | gal x long | Later-generation |
| EIC_11 | 0.274 | 0.148 | gal | gal x long | Later-generation |
| SE_01_03 | 0.866 | 0.091 | gal | gal x long | Backcross gal |
| SE_01_04 | 0.763 | 0.346 | gal | gal x long | Backcross gal |
| SE_12_04 | 0.547 | 0.187 | gal | gal x long | Later-generation |

Comparing genotypes from the parental species *D. galeata* and *D. cucullata* (five and three individuals, respectively) led to identifying 715,438 fixed sites between the two species (due to quality filtering, 275,216 of these sites were further analyzed). All three *D. galeata* x *D. cucullata* hybrids carried a *D. cucullata* mitochondrial genome, their hybrid index varied between 0.079 and 0.267 and their heterozygosity ranged from 6.4% to 50.9 % (Figure 2B&C, Table 2). The individual SE_23_04 is most likely the result of a backcrossing with *D. galeata*; however, it is difficult to determine what backcrossed with it: either an F1 hybrid or a later generation hybrid i.e., that resulted from several generations of admixture. Haplotype information would be needed to gain certainty. The other two hybrids’ lower hybrid index hints backcrossing with *D. galeata,* according to the criteria defined above.

### Population genomics parameters

To calculate genome-wide nucleotide diversity (π), between-taxon differentiation (F_ST_), and between-taxon divergence (d_xy_) within 100-kb sliding windows, we took advantage of the inference made with ADMIXTURE and pooled all genotypes which were unambiguously assigned to either of the parental species clusters. Consequently, a total of seven genotypes from four populations were classified as *D. cucullata*, eight from five populations as *D. longispina*, and 20 genotypes from eight populations as *D. galeata* (Table S2). All values (d_xy_, π and F_ST_) were calculated with the script popgenWindows.py (github.com/simonhmartin/genomics_general release 0.3) for each species pair and are plotted for the 50 largest scaffolds in Figure 3.

**Figure 2 Figure 3:**
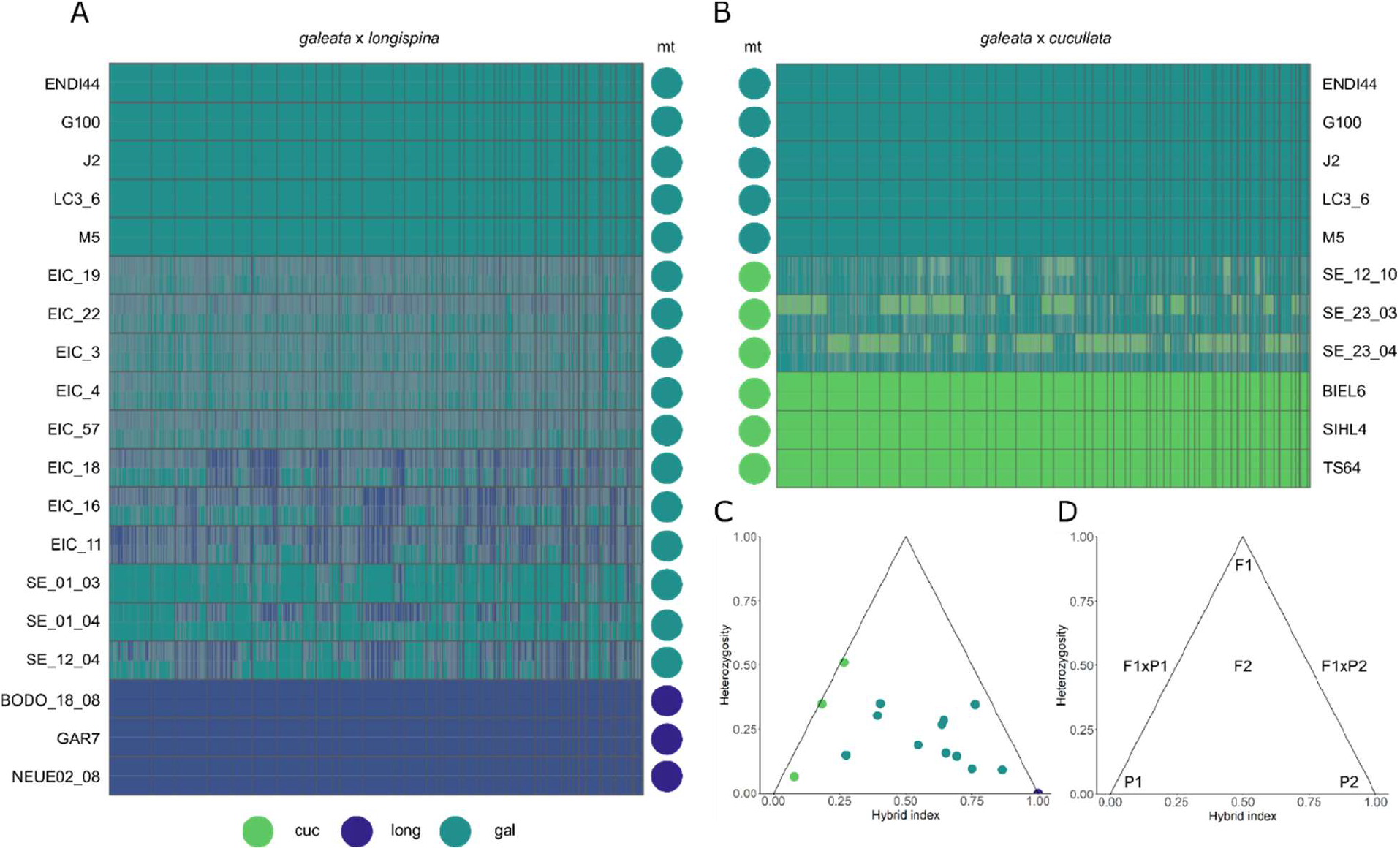
**Panels A & B** Ancestry painting of the hybrid individuals identified through the admixture analysis. Each row represents an individual. Colored circles on the side indicate the mitochondrial identity of the individuals, based on the analysis of full mitochondrial genomes. Scaffolds are sorted by length and separated by thin grey lines. In panels **A** and **B**, the five upper rows represent individuals assigned to the parental species *D. galeata*. In **A,** the last three rows correspond to individuals assigned to the parental species *D. longispina.* In **B.** the last three rows correspond to individuals assigned to the parental species *D. cucullata* Triangle plots summarizing **C.** the hybrid index and mitochondrial species identity for all individuals identified as admixed **D.** the hypothetical expected means of parental species (P1 and P2) and hybrid classes (F1xP1 and F1xP2: backcrosses with parental species P1 and P2, respectively).

The window-based F_ST_ values for all three possible pairs among the three species averaged 0.274 for *D. galeata* vs *D. longispina*, 0.343 for *D. cucullata* vs *D. longispina* and 0.364 for *D. galeata* vs *D. cucullata*. The mean sequence divergence d_xy_ for the three pairs was 0.018 for *D. galeata* vs *D. longispina* and 0.022 for both *D. cucullata* vs *D. longispina* and *D. cucullata* vs *D. galeata*. Both parameters show similar patterns, with lower values on average when comparing *D. galeata* to *D. longispina* than when comparing *cucullata* to either one of the other species. These patterns confirm the results obtained with other analyses, for example, the higher number of fixed sites between *D. galeata* and *D. cucullata* in the ancestry painting analysis.

The window-based estimates show high variability in levels of differentiation and divergence along the genome. Further, regions of high or low differentiation are mostly associated with depleted or high nucleotide diversity, respectively (see scaffolds 2 and 9 for example). However, the genome being represented by unordered scaffolds instead of chromosomes makes this difficult to interpret further.

Nucleotide diversity (π) to quantify the level of genetic variation within each taxon was on average higher for *D. longispina* (1.18%) than for the other two species (0.95% and 0.85% for *D. galeata* and *D. cucullata*, respectively). This cannot be explained by the differences in group sample sizes, since *D. galeata* was the group with the largest sample size (and highest number of sampled populations). To ensure our window-based estimates were not biased because of the overrepresentation of some populations in a group (e.g. DOB in the *galeata* group), we also calculated these indices using only one individual from each population per species; if one population contained multiple individuals, we picked one individual at random to represent this population (see Table S2 for a listing of the used genotypes - results shown in Figure S4).

Many more highly differentiated windows and genes were shared among two or all species pairs than would be expected by a random intersection (Figure S5). For example, a total of 2575 10kb windows had an FST value within the 95^th^ percentile in the pair *D. galeata*/ *D. longispina* and 2569 in the pair *D. galeata*/ *D. cucullata*. The mean expected number of windows in common between these two pairwise comparisons was 113, but the number of windows in common observed in the data was 1601. A similar pattern was observed in all other intersections. This result suggests that the location of differentiated genome parts is not due to random processes but has biological significance. A GO-enrichment analysis of these isolated genes to shed light on the function of these species-specific genes, however, was not possible, because of the low number of genes with GO annotation. For the pair *D. galeata/cucullata*, only 12% of the genes in the outlier windows were annotated with Gene Ontologies, for the pair *D. galeata/longispina* it was 11% and for the *D. cucullata/longispina* pair it was 10%.

### Phylogeny based on complete mitochondrial genomes

Phylogenetic reconstruction based on the mitochondrial protein-coding and ribosomal RNA genes were largely consistent with earlier mitochondrial phylogenies based on single or few mitochondrial genes (e.g. Adamowicz*, et al.* 2009; Petrusek*, et al.* 2012). We identified highly supported clades comprising the respective parental genotypes, hence representing *D. longispina*, *D. cucullata*, and *D. galeata* mitochondrial haplotypes (Figure 4). *D. cucullata* and *D. galeata* mitochondrial haplotypes clustered as sister groups. While the mitochondrial haplotypes in the *D. longispina* and *D. cucullata* clusters do not show much divergence, the *D. galeata* haplotype cluster also contains deeper branching events (haplotype EIC_15 and AR3_17 / AR5_18). Further, although all samples from AR were unequivocally categorized as *D. cucullata* in the ADMIXTURE analysis and clustered with *D. cucullata* parental genotypes in the PCA, two of them have mitochondrial haplotypes falling into the *D. galeata* cluster (AR3_17 and AR5_18). A similar mismatch was also observed for EIC_15, which clusters with *D. longispina* when considering nuclear SNP and with *D. galeata* when considering the mitochondrial genome. Within the species clusters, we observed a grouping by lake with many haplotypes being either identical or very similar when originating from the same location. The trees obtained with either only protein-coding genes (CODON model) or protein-coding and ribosomal RNA genes but with a mixed model (DNA for rRNAs and CODON for DNA) were all consistent with the tree shown here and are therefore not included.

**Figure 4:**
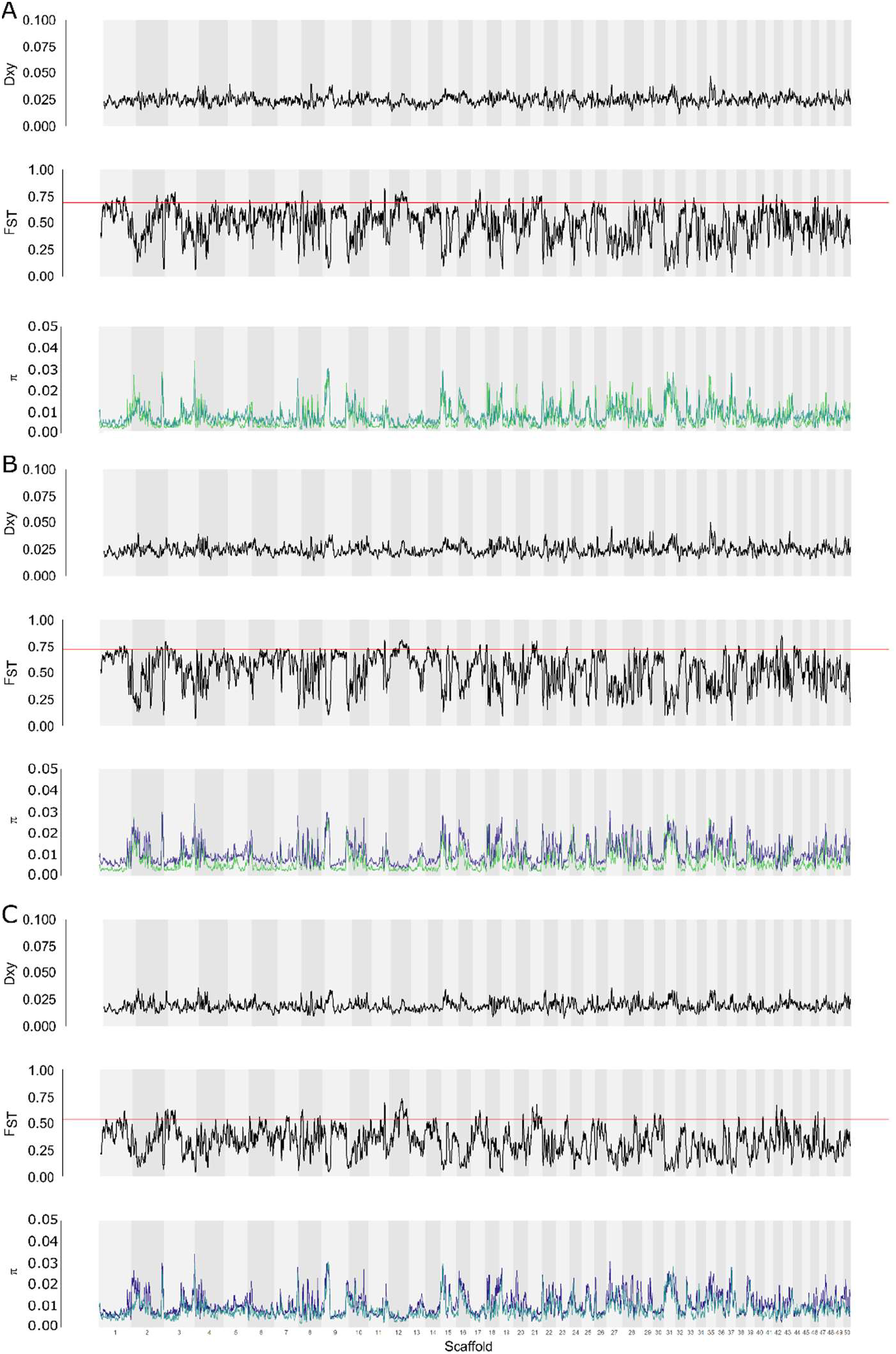
Window-based statistics for the pairs **A**. *D. galeata* / *D. cucullata,* **B**. *D cucullata* / *D. longispina* and **C**. *D. galeata* / *D. longispina*, shown for the 50 largest scaffolds in 100kb windows with 10kb step size – calculations are for all non-admixed individuals unambiguously assigned to parental species according to the ADMIXTURE analysis. **In each panel from top to bottom:** d_xy_ values, pairwise F_ST_ values with a red horizontal line indicating the 95^th^ percentile, nucleotide diversity (π) for *D. galeata* (teal), *D. longispina* (dark blue), and *D. cucullata* (lime green).

### Patterns of introgression

We tested all four northern Germany populations (EIC, DOB, AR and SE) for admixture between the three reference species with *f*_3_ statistics tests (Table 3) and considered a Z-score < −3 as significant (following Patterson*, et al.* 2012; Reich*, et al.* 2009). Negative and significant values (*f*_3_=−0.19) using EIC as the test population and *D. galeata* and *D. longispina* as the source populations indicated mixed ancestry from these two or closely related populations. For population SE, the *f*_3_ test supports both admixed ancestry from *D. galeata* and *D. longispina* (*f*_3_=-0.09) and *D. galeata* and *D. cucullata* (*f*_3_=-0.15). All tests for population DOB and AR were positive providing no evidence of admixture events. The supported introgression events are consistent with the results in our previous analyses conducted with ADMIXTURE and the ancestry painting approach.

**Table 3:** Summary of the *f*_3_ statistic for admixture in the form (C; A, B). A significantly (Z-score < −3, in bold) negative *f*_3_ value implies that the target population C is admixed. SE: standard error

| Source population A | Source population B | Target population C | $f_3$ | SE | Z-score | # Sites |
| --- | --- | --- | --- | --- | --- | --- |
| gal | long | AR | 5.48226 | 0.154392 | 35.509 | 1,072,106 |
| gal | cuc | AR | 0.245124 | 0.006208 | 39.487 | 791,570 |
| long | cuc | AR | 0.225393 | 0.005475 | 41.165 | 875,293 |
| gal | long | Dob | 0.152725 | 0.003233 | 47.242 | 752,265 |
| gal | cuc | Dob | 0.150027 | 0.003028 | 49.546 | 815,035 |
| long | cuc | Dob | 1.763147 | 0.049657 | 35.506 | 1,059,418 |
| gal | long | EIC | -0.193114 | 0.003224 | **-59.898** | 864,328 |
| gal | cuc | EIC | 0.086614 | 0.002623 | 33.025 | 1,119,715 |
| long | cuc | EIC | 0.014322 | 0.001774 | 8.071 | 1,145,530 |
| gal | long | SE | -0.093233 | 0.003222 | **-28.939** | 1,029,744 |
| gal | cuc | SE | -0.154696 | 0.003924 | **-39.426** | 955,516 |
| long | cuc | SE | 0.288212 | 0.011203 | 25.727 | 1,092,186 |

We performed local ancestry inference with Loter (Dias-Alves*, et al.* 2018) to trace genome-wide introgression among the hybrids and infer additional details about the parental species and backcross history from haplotype information. The results were summarized genome-wide for the ancestry proportion, heterozygosity of ancestry and the number of ancestry transitions where each ancestry tract is counted when the state of an SNP changes to the other species or at the end of a scaffold (Figure 5). The three *D. galeata* x *cucullata* hybrids were all found to have high *galeata* ancestry (73.9%-94.2%) and the individuals SE_23_03 and SE_23_04 were confirmed as the offspring of a later-generation hybrid and a pure *galeata* parent (Figure S6B).

**Figure 5:**
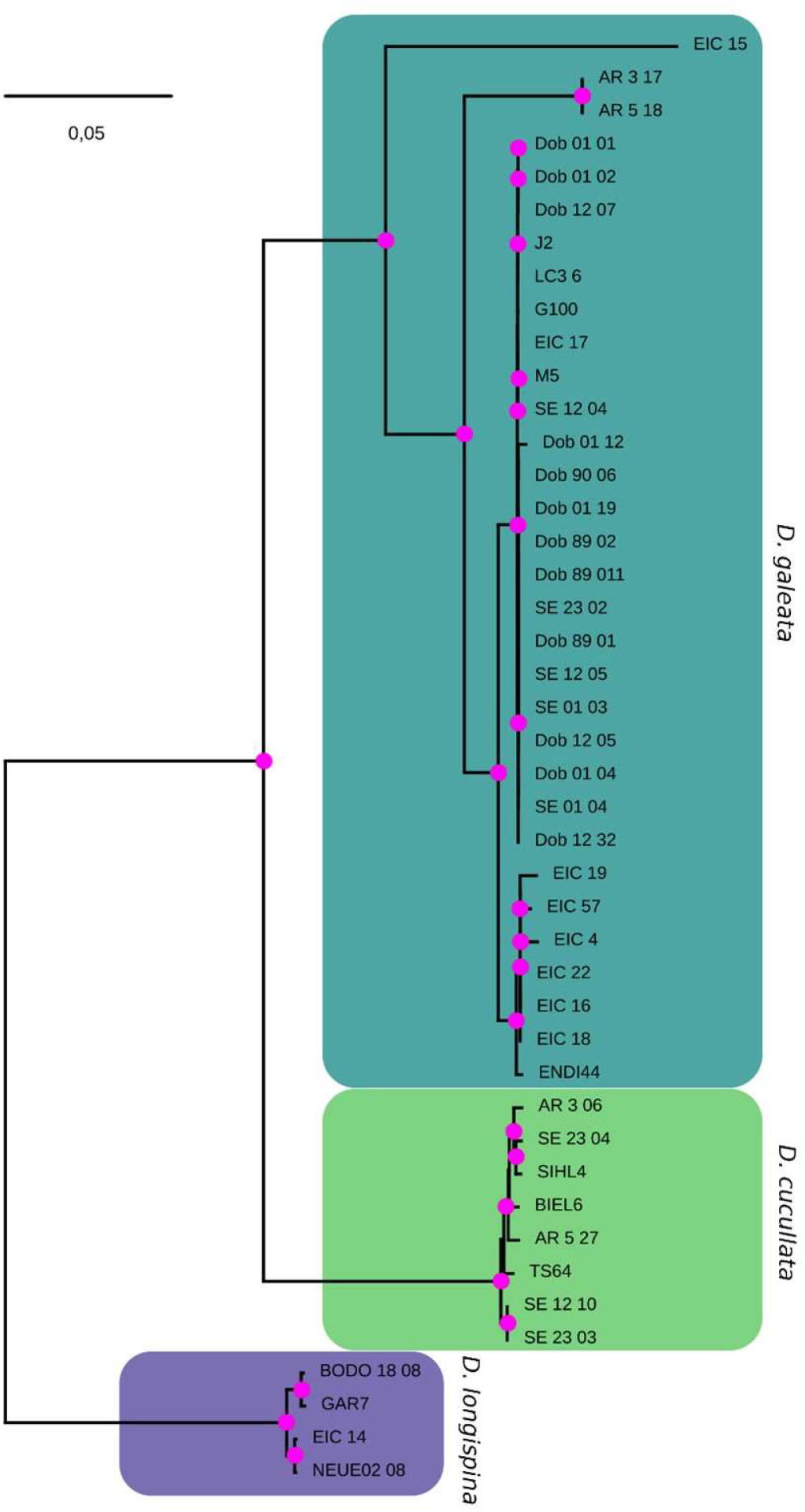
Maximum-likelihood tree reconstructed from mitochondrial protein-coding and ribosomal RNA genes of parental species, clones sampled in the water column and resting eggs sequenced in this study. The tree reveals distinct and highly supported clusters corresponding to *D. galeata*, *D. cucullata* and *D. longispina* mitotypes (as defined by the respective parental species and a sister taxa relationship between *D. galeata* and *D. cucullata)*. Here, the best tree (logL = −47950.82) rooted with outgroup *D. laevis* is depicted. Magenta dots indicate Shimodaira-Hasegawa approximate likelihood ratio test values >= 80% and ultrafast bootstrap support values >= 95% calculated from 10,000 bootstrap replicates (SH-aLRT / UFboot). The scale bar corresponds to 0.05 nucleotide substitutions per nucleotide site. KL11 was excluded due to missing data.

Seven *D. galeata* x *longispina* hybrids had very high *galeata* ancestry (81.4%-97.9%), and visual inspection of the ancestry tracts (Figure S6A) revealed very short *longispina* tracts and scaffolds with multiple breakpoints indicating multiple generations of recombination. Four *D. galeata* x *longispina* hybrids had lower *galeata* ancestry (27.7%-59.0%) and the presence of complete *longispina* scaffolds implying some backcrossing with *longispina*. The haplotype phasing confirmed that the parents of all *D. galeata* x *longispina* hybrids were also of hybrid origin. The average and maximum ancestry tract length for all *D. galeata* x *longispina* hybrids is shorter than those for *D. galeata* x *cucullata* hybrids.

**Figure 6:**
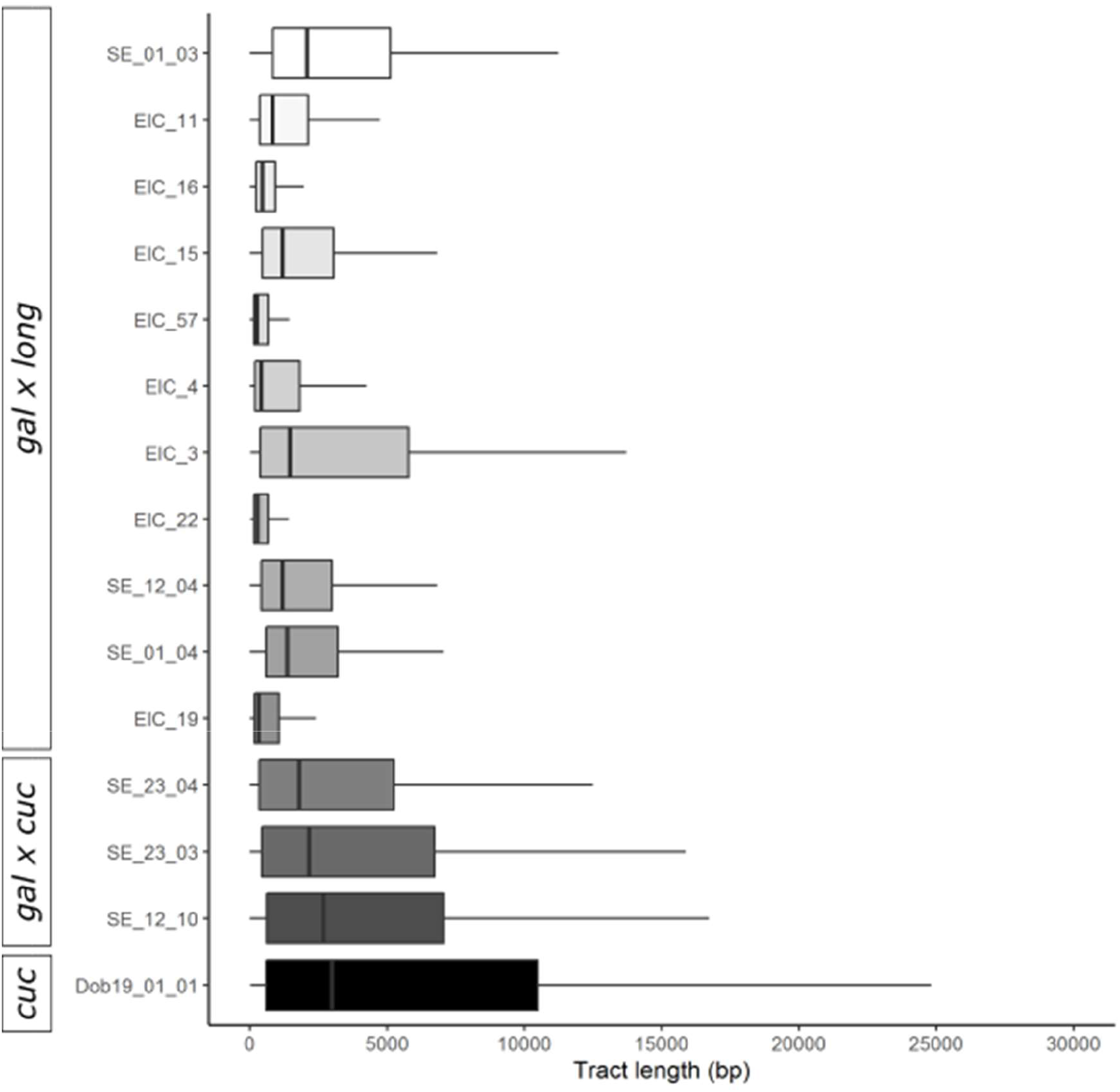
Distribution of the ancestry tract length where each ancestry tract represents the state of a SNP changing to the other species or the end of a scaffold in the local ancestry inference for each admixed individual and one non-admixed *D. cucullata* individual. The non-admixed individual displays the ancestry tract length distribution when all scaffolds derive from the same species. Hybrid type (according to ADMIXTURE analysis) is given on the left side.

## Discussion

### A reference genome for studying a species complex

*Daphnia* are a key species in freshwater habitats. Previous studies have established reference genomes for the model species *D. pulex* (Colbourne*, et al.* 2011; Ye*, et al.* 2017) and *D. magna* (Lee*, et al.* 2019). No high-quality reference genome for species belonging to the *Daphnia longispina* species complex was available so far. To date, it is unclear whether the ecological differenciation and/or intrinsic incompatibilities drive and maintain divergence between DLSC species. Besides its utility for studies of hybridization events in the DLSC, the new assembly we present here will thus allow us to better understand the evolution of a key species in European freshwaters.

Even though the onset of the DLSC radiation was dated to 5-7 Mya based on nuclear and mitochondrial markers (Adamowicz*, et al.* 2009; Schwenk 1993; Taylor*, et al.* 1996), several factors confirm the suitability of this reference for all tested species. Mapping success and coverage of whole-genome data from *D. cucullata* and *D. longispina* to the reference genome were high, and we found no evidence of reference bias. This assembly clearly benefited from advances both in the sequencing technologies and assembly and post-processing algorithms since the first *Daphnia* genome (Colbourne*, et al.* 2011). The metrics used for assessing its quality reveal that in particular, the combination of long and short read technologies led to highly contiguous and accurate scaffolds. Although we likely did not recover the genome in its full length (133Mb out of an estimated 156Mb), and the N50 value is lower than those obtained for *D. pulex* (Ye*, et al.* 2017) and *D. magna* (Lee*, et al.* 2019), iterative scaffolding allowed for a very efficient gap-closing, and an exceptionally low number of mismatches, compared to the other *Daphnia* assemblies.

### Pervasive introgression in the *Daphnia longispina* species complex

We utilize a method that allows us to interrogate biological archives and analyze whole *Daphnia* genomes directly from the resting egg bank (Lack*, et al.* 2018) without hatching and culturing several clonal lineages. This provides a wide sweep of populations, past and present, with each egg being the product of local sexual recombination.

While no evidence of introgression was found in the DOB population, the three other locations host a variety of admixed genotypes. SE & EIC can even be considered hybridization hotspots with more than 60% of individuals having hybrid ancestry, as revealed in the ADMIXTURE analysis. However, Kong and Kubatko (2021) very recently showed that ADMIXTURE is sensitive to unequal contributions by the parental species, and we thus sought to support these inferences by *f*_*3*_ calculations and using an ancestry painting approach.

In DOB, ADMIXTURE delivered unequivocal results. Further, the *f*_*3*_ index indicated that no introgression was detectable in this population. However, the PCA plot shows that some of the DOB genotypes are near hybrid individuals. ANGSD results are similar but these genotypes nearer the parental species (Figure S3A). To address these slightly contradictory results, we therefore conducted an ancestry painting on two DOB genotypes, (12_07 and 89_02, Table S7). Both genotypes had a very low heterozygosity, thus confirming the ADMIXTURE and *f*_*3*_ outcomes. A possible explanation would be that these two genotypes carry variation that is not reflected in our limited sampling of the parental species. When comparing to microsatellite-based analysis including many more populations and data points (e.g. Thielsch*, et al.* 2009), the *D. galeata* cluster has often been larger and more diverse than the others. The seemingly two “stray” DOB genotypes are therefore likely well within the species variation boundaries. All mitochondrial haplotypes were clustered together in the phylogenetic reconstruction as well.

In AR, despite the high resting egg density found in the sediment, only very few could be successfully genotyped. While all inferences based on nuclear markers (PCA, ADMIXTURE, *f*_*3*_) indicated an absence of hybridization or introgression in this population, the mitochondrial phylogenetic reconstruction showed diverging results. From a nuclear point of view, all genotypes could be categorized as *D. cucullata*, but two out of four AR individuals presented the mitochondrial genome of another species, i.e., *D. galeata*. However, the phylogenetic reconstruction shows that the two AR mitochondrial haplotypes form a cluster separate from the main *D. galeata* cluster, which hints at different evolutionary history for these mitochondrial genomes. Such distinct lineages within a species and mito-nuclear discordances were also found by Thielsch*, et al.* (2017) in the DLSC and mitochondrial capture has been detected in other *Daphnia* species (Marková*, et al.* 2013). It is an interesting phenomenon in the DLSC that merits to be further investigated in the future with broader sampling.

In EIC, both pelagic samples and resting eggs were analyzed. Genotypes sampled alive from the water column were all inferred to be admixed to various degrees, three resting egg samples were also admixed, and the remaining five were assigned to either one of the 2 parental species. We conducted the ancestry painting approach on all EIC individuals; the fixed sites heterozygosity of the individuals categorized as non-admixed in ADMIXTURE was indeed near zero (Table S7). Such high abundances of *D. galeata* x *longispina* hybrid resting eggs in periods of rapidly changing environmental conditions (i.e. eutrophication) have also been recorded in Lake Constance (Brede*, et al.* 2009). The high frequency of *D. galeata* x *longispina* hybrids observed here might be due to similar ecological history: the lake Eichbaumsee was created through sand excavation for construction work around ~40 years ago and is characterized by extreme eutrophication and even hypertrophy that could not be remediated. The presence of later-generation hybrids and backcrosses with *D. galeata* and *D. longispina* and short ancestry tract length suggest that both species have been present and hybridizing for most of the lake’s short history, or even that it was colonized by individuals of hybrid origin. However, we only obtained contemporary samples for EIC and analysis of resting eggs from sediment cores are needed to distinguish between the two hypotheses.

In SE, diversity is high, both in terms of species combinations in admixed individuals and in terms of degrees of introgression. Although we analyzed eggs from sediments cores, they originated from the first centimeters and there is, therefore, no clear temporal pattern that separates the different hybrid combinations found here. Strikingly, while SE and DOB are geographically very close to each other (~10 km), and dispersal of resting eggs through e.g., waterfowl or storms would be possible (Figuerola*, et al.* 2005; Frisch*, et al.* 2007; Pietrzak and Slusarczyk 2006), the *Daphnia* communities are quite different. This might be due to their different eutrophication levels, reflect the fact that initial colonization was followed by the establishment of different species, or a combination of both. The observed diversity at such a small spatial scale underlines the mosaic nature of freshwater habitats and the usefulness of approaches including many populations to fully understand genetic diversity arising from colonization and hybridization events in the DLSC.

Previous studies using mitochondrial and few nuclear markers (e.g. Alric*, et al.* 2016; Thielsch*, et al.* 2012) were able to categorize hybrids into F1, F2 and backcrosses. However, due to the low resolution of the used markers, further categorizing and above all identification of genome-wide breaking points was not possible at the time. The *D. galeata* reference genome and resequencing data offer now a much higher resolution to assess later generation hybrids and patterns across the genome. In general, hybrids identified in this study seem to have a complex history of multiple generations of hybridization and backcrossing with both parental species that we are not able to detangle using only ancestry paintings. The local ancestry inference revealed that the average ancestry tract length for *D. galeata* x *longispina* hybrids from EIC and SE is shorter than those for *D. galeata* x *cucullata* hybrids. There are several explanations for the observed pattern. One is that more generations of recombination led to shorter introgressed tracts, and the *D. galeata* x *longispina* hybrids are therefore the result of a greater number of sexual generations than the *D. galeata* x *cucullata* hybrids. The genomic mosaic of ancestry segments for all hybrid individuals is also characterized by multiple breakpoints within the same scaffolds, which is only possible after multiple generations of recombination. However, data on genome-wide recombination rates and selection are needed to reach solid conclusions about the correlation between tract length and age of the hybridization event in the individual’s ancestors. Alternatively, reproductive isolation might be lower between *D. galeata* and *D. longispina* than between *D. galeata* and *D. cucullata*, thus leading to faster introgression in the former case.

As evidenced by the comparison of genomic windows of higher divergence between species pairs, the introgression pattern is not random: a given region exhibiting high FST values between the *D. galeata* and *D. longispina* genotypes is also likely to show similarly high FST values in the *D. galeata*/ *D. cucullata* pair. Further, some parts of the genome seem to be effectively shielded from introgression. About a quarter of all genes (4136) are in regions that are highly differentiated between at least two species and about 5% (859) in parts of the genome that are isolated among all three species of the complex. This is much more than expected by chance (Figure S3) and is thus likely due to selection against introgression. It seems plausible to search among these for genes that conserve the specific identity of the involved taxa, despite incomplete reproductive isolation. Genes responsible for the observed ecological divergence among the taxa (Schwenk*, et al.* 2000) or genetic incompatibilities are most likely candidates to be found in the observed divergent regions. Given the ancient divergence, the speciation process in the DLSC might have attained a selection-migration-drift equilibrium, for which there is growing empirical evidence in other species like stick insects (Riesch*, et al.* 2017), flycatchers (Burri*, et al.* 2015), and non-biting midges (Schreiber and Pfenninger 2020). However, the current snapshot could equally likely be a consequence of one or several pulses of hybridization. To assess the stability of the equilibrium, data showing that the introgression/selection process is ongoing and constant across an extended period of time would be required and *Daphnia* offers the unique opportunity to go back in time to test these alternative hypotheses.

### New evidence for cytonuclear discordance

The genome-wide perspective also elucidated discordance between nuclear and mitochondrial patterns. The phylogeny based on mitochondrial genomes conforms to previously inferred relationships in the DLSC and suggests *D. galeata* and *D. cucullata* are sister species, with *D. longispina* as an outgroup (Adamowicz*, et al.* 2009; Petrusek*, et al.* 2012). However, several of our analyses based on nuclear SNPs challenge this view and suggest different evolutionary histories for mitochondrial and nuclear genomes. The ancestry painting approach relies on the identification of fixed sites for species in a pairwise comparison. More sites were found to be fixed between *D. galeata* and *D. cucullata* (715,438) than between *D. galeata* and *D. longispina* (335,052), which implies a greater divergence between members of the former pair. Further, FST values were on average higher between *D. galeata* and *D. longispina* (0.274) than between *D. galeata* and *D. cucullata* (0.364). Reports of cytonuclear discordance are common both in plants (e.g. Folk*, et al.* 2017; Huang*, et al.* 2014; Lee‐Yaw*, et al.* 2019; Stephens*, et al.* 2015) and animals (e.g. Llopart*, et al.* 2014; Melo-Ferreira*, et al.* 2014; Sarver*, et al.* 2021). Several processes can lead to this discordance among closely related species: incomplete lineage sorting causing phylogenetic reconstructions based on mitochondrial markers to differ from the true phylogeny of the taxa, or selection causing the fixation of different mitochondrial genomes in different places from standing variation within species (e.g. Barrett and Schluter 2008). Alternatively, cytonuclear discordance may reflect hybridization between species and cytoplasmic introgression, accompanied or not by selection (reviewed in Sloan*, et al.* 2017). The latter explanation would be quite conceivable in the DLSC.

## Conclusion

We here provide the first high-quality resources to study genome-wide patterns of divergence in the *Daphnia longispina* species complex, an ecologically important taxon in European freshwater habitats. By quantifying intra- and interspecific diversity, we provide a first glimpse into introgressive hybridization and lay the ground for further studies aiming at understanding how species boundaries are maintained in the face of gene flow.

Unlike for *D. pulex* and *D. magna*, no linkage groups are known for any species of the DLSC. Hi-C sequencing data will be added in the future to order scaffolds into larger, potentially chromosome-scale scaffolds. Such an approach holds promise in a species complex were laboratory crossings for F2 panels and traditional mapping are nearly impossible. This will allow discovering structural variants, identifying recombination breakpoints along each chromosome and thus provide a deeper understanding of the introgression patterns observed here. The functional role of genes in the regions of high divergence uncovered through this first analysis is yet unclear and will be addressed in future studies.

Finally, wider sampling, with the inclusion of more populations as well as more members of the species complex, and the reconstruction of a nuclear based phylogeny are necessary to reach conclusions about the species relationships and eventually identify the causes of the pattern uncovered here.

## Materials & Methods

### Sampling

The clonal line used for genome sequencing and assembly, M5, was hatched from a resting egg isolated from the upper layers (first 5cm, corresponding to the years 2000-2010) of a sediment core taken in Lake Müggelsee in 2010. Further, single genotypes representing the parental species from various locations were used in this study, henceforth “parental species genotypes”. Most of them were established from individuals sampled from the water column and are still maintained through asexual reproduction as monoclonal cultures in the laboratory. Thus, all individuals of a clonal line are the same genotype and can be pooled to achieve sufficient amounts of genomic DNA. The species identity for these genotypes was established through a combination of methods: morphology, mitochondrial sequences, and nuclear markers.

Sediment cores were collected from Dobersdorfer See (DOB), Selenter See (SE) and Arendsee (AR), Germany using a gravity corer (Uwitec, Mondsee, AT) (Table S4). Samples were taken from the deepest part of the lakes to minimize past disturbance of the sediment. Cores were cut horizontally into 1cm layers and the layers were stored at 4 °C in the dark to prevent hatching. Sediment rate of the three lakes was determined using radioisotope dating (^137^Cs and ^210^Pb).

In addition, lake sediment from the shoreline of Eichbaumsee (EIC) (Table S4), Germany was collected by hand and stored at 4 °C. The exact age of the sediment is unknown but the upper layers most likely contain recent eggs from the last few years. Zooplankton samples were taken from Eichbaumsee using a plankton net (mesh size 150 μm) from which six *Daphnia* clonal lines were established in a laboratory setting with artificial medium (Aachener Daphnien Medium, ADaM Klüttgen*, et al.* 1994).

All sampling locations are plotted in Figure 1A and information on all samples is provided in Table S2.

### Genome sequencing

#### DNA extraction for genome sequencing with Illumina & PacBio

DNA was extracted from around 60 clonal M5 individuals collected from batch cultures maintained in ADaM, and fed with the algae *Acutodesmus obliquus*, cultivated in medium modified after (Zehnder and Gorham 1960). Extraction was conducted following a phenol chloroform-based protocol with an RNase step and subsequently sequenced on an Illumina HiSeq4000 at BGI China. Additionally, tissue samples with around 3000 individuals were sent to BGI for DNA extraction and PacBio sequencing.

#### Re-sequencing (Population genomics approach)

##### DNA extraction from batch cultures for re-sequencing

For clonal lines used as reference for the parental species, individuals were raised in batch cultures and treated with antibiotics prior to collection and storage at −20 or −80°C. DNA was extracted with either a phenol chloroform method, a (modified) CTAB protocol or a rapid desalting method (MasterPure™ Complete DNA and RNA Purification Kit; Lucigen Corporation).

Total genomic DNA was isolated from 20 pooled adult *Daphnia* for each of the five EIC clonal lines using a CTAB extraction method (Doyle and Doyle 1987).

##### Whole Genome amplification on resting eggs for re-sequencing

To isolate *Daphnia* resting eggs from the sediment each sediment layer was sieved using a sieve with 125 μm mesh size and small amounts of the remaining sediment were resuspended in distilled water. Ephippia were eye spotted under a stereomicroscope, counted and transferred to 1.5 mL tubes. The water was removed and ephippia stored at −20 °C in the dark until further analysis.

The ephippia were then opened under a binocular with insect needles and tweezers previously treated under a clean bench (UV sterilization) and with DNase away (Thermo Fisher). Eggs that were already damaged, had an uneven shape or were orange, which is evidence for degradation, were discarded. The resting egg separated from the ephippial casing washed in 15 μl sterile 1x PBS and then transferred in 1 μl 1x PBS to a new tube with 2 μl fresh 1x PBS. The isolated eggs were stored at −80 °C at least overnight.

For whole genome amplification of single eggs, the REPLI-g Mini Kit (Qiagen) was used. This kit is enabling unbiased amplification of genomic loci via Multiple Displacement Amplification (MDA). The isolated resting eggs were thawed on ice and the whole genome was amplified following the manufacturer’s protocol for amplification of genomic DNA from blood or cells. Briefly, denaturation buffer was added to the prepared resting eggs in 3 μl 1x PBS and amplified by REPLI-g Mini DNA Polymerase under isothermal conditions for 16 hours.

The amplified product was quantified on a Nanodrop spectrophotometer (Thermo Fisher) and with a Qubit Fluorometer (Thermo Fisher). Successful amplifications were purified with 0.4 x Agencourt AMPure XP magnetic beads (Beckman Coulter) to remove small fragments and eluted in 60 μl 1x TE buffer.

Fragments of the mitochondrial gene 16S rRNA gene were amplified to check successful amplification of *Daphnia* DNA using the universal cladoceran primers S1 and S2 (Schwenk*, et al.* 1998) and a low presence of bacterial DNA using universal primers for the bacterial 16S rDNA gene (Nadkarni*, et al.* 2002). Only samples with a successful amplification of the *Daphnia* 16S fragment and low amplification of the bacterial 16S fragment indicating low bacterial contamination were used for sequencing steps.

##### Library preparation and sequencing of re-sequencing samples

After quantification and quality control of the DNA using Nanodrop and Qubit instruments, libraries were prepared either directly in-house with the NEBNext^®^ Ultra™ II DNA Library Prep Kit for Illumina^®^ (New England Biolabs), or at the sequencing company Novogene (Cambridge, UK). Resequencing (paired-end 150bp reads) was then performed either at Novogene (UK) Company Limited or the Functional Genomics Center (ETH Zurich and University of Zurich) on Illumina NovaSeq 6000 and HiSeq4000 instruments.

Details on the procedure used for each sample are provided in Table S2.

#### Genome assembly and annotation

We provide here a summarized version of the procedure used to assemble and annotate the genome. Details can be found in Supplementary Methods.

##### Raw data QC

Illumina reads were trimmed and the adapter removed using a combination of Trimmomatic 0.38 (Bolger*, et al.* 2014), FastQC 0.11.7 (Andrews 2010) and MultiQC 1.6 (Ewels*, et al.* 2016) within autotrim 0.6.1 (Waldvogel*, et al.* 2018). To filter out reads possibly originating from contamination from known sources (see below), a FastQ Screen like approach was chosen. In brief, the reads are separated by results of mapping behavior to different genomes. Positive controls consisted of genome data for other *Daphnia* species (dmagna-v2.4 and Daphnia_pulex_PA42_v3.0, see Supplementary Methods for accession numbers), and negative control i.e. sequences deemed undesirable for genome assembly consisted of genome data from human, bacteria, viruses and the algae used to feed the batch cultures. The resulting database comprised 108,163 sequences (total sequence space 42.2 Gb). Both Illumina reads and PacBio subreads were mapped against the database with NextGenMap (Sedlazeck*, et al.* 2013) and minimap2 (Li 2018), respectively.

Reads did only pass the filtering if they either did not map to the database at all or had at least one hit against one of the two *Daphnia* genomes. Table 2 in Supplementary methods gives an overview of the effect of different filtering steps.

##### Assembly and contamination screening

All paired and unpaired contamination filtered Illumina reads as well as the contamination filtered PacBio reads were used as input for RA 0.2.1 (https://github.com/rvaser/ra). Blobtools 1.0 (Laetsch and Blaxter 2017) was used to screen the resulting assembly for possible unidentified contamination in the hybrid assembly. Briefly, bwa mem 0.7.17 (Li 2013) was used to map Illumina reads back to the assembly and taxonomic assignment was done by sequence similarity search with blastn 2.9.0+ (Camacho*, et al.* 2009). Contamination with different bacteria was clearly identifiable, and contigs with coverage below 10x and/or GC content above 50% were removed. Additionally, PacBio reads mapping to these contigs were removed to minimize false scaffolding in further steps. The contig corresponding to the mitochondrial genome was identified after a blast search against available mitochondrial genomes for this species and removed from the assembly.

##### Scaffolding and gap closing

The blobtools filtered PacBio reads were used for scaffolding and gapclosing, which was conducted in three iterations. Each iteration consisted of a scaffolding step with SSPACE LongRead 1-1 (Boetzer and Pirovano 2014), a gap closing step with LR Gapcloser (https://github.com/CAFS-bioinformatics/LR_Gapcloser; commit 156381a), and a step to polish former gap parts with short reads using bwa mem 0.7.17-r1188 and Pilon 1.23 (Walker*, et al.* 2014) in a pipeline developed to this effect, wtdbg2-racon-pilon.pl 0.4 (https://github.com/schellt/wtdbg2-racon-pilon).

##### Assembly quality assessment

Contiguity was analyzed with Quast 5.0.2 (Gurevich*, et al.* 2013) at different stages of the assembly process. Further, mapping rate, coverage and insert size distribution were assessed by mapping Illumina and PacBio reads with bwa mem and Minimap 2.17 respectively. To show absence of contamination in the assembly blobtools was ran as above. The genome size was estimated by dividing the mapped nucleotides by the mode of the coverage distribution of the Illumina reads by backmap 0.3 (https://github.com/schellt/backmap), resulting in 156.86Mb (with the obtained assembly length amounting to 85% of this estimated length). Additionally, the genome size was estimated using a k-mer based approach by creating a histogram from raw Illumina reads with Jellyfish 1.1.12 (Marçais and Kingsford 2011) and running the GenomeScope web application (http://qb.cshl.edu/genomescope/) resulting in a genome size estimate of 150.6Mb.

Completeness in terms of single copy core orthologs of the final scaffolds was assessed with BUSCO 3.0.2 (Simão*, et al.* 2015), using the Arthropoda set (odb9).

##### Genome Annotation

RepeatModeler 2.0 (Smit and Hubley 2015) was run to identify *D. galeata* specific repeats. The 1,115 obtained repeat families were combined with 237 *D. pulex* and 1 *D. pulicaria* repeat sequences from RepBase release 20181026 to create the final repeat library. The genome assembly was then soft masked with RepeatMasker 4.1.0 (Smit*, et al.* 2013-2015), resulting in 21.9% of the assembly being masked.

Gene prediction models were produced with Augustus 3.3.2 (Stanke*, et al.* 2008), GeneMark ET 4.48_3.60_lic (Lomsadze*, et al.* 2005) and SNAP 2006-07-28 (Korf 2004). The Augustus model was based on the soft masked assembly and the *D. galeata* transcriptome (HAFN01.1, Huylmans*, et al.* 2016). The GeneMark model was obtained by first mapping trimmed RNAseq reads to the assembly with HISAT 2.1.0 (Kim*, et al.* 2019) and then processing the resulting bam file with bam2hints and filterIntronsFindStrand.pl from Augustus to create a gff file with possible introns, which was finally fed into GeneMark.

The structural annotation was conducted in MAKER 2.31.10 (Holt and Yandell 2011). Briefly, the unmasked genome assembly, the species own transcriptome assembly as ESTs, the complete Swiss-Prot 2019_10 (UniProt Consortium, 2019) and the protein sequences resulting from *D. magna* (Lee*, et al.* 2019), as well as *D. pulex* (Ye*, et al.* 2017) genome annotations as protein evidence, were used as input for MAKER. In total three iterations of MAKER with retraining of the Augustus and SNAP model in between the iterations were conducted.

The quality of the structural annotation was assessed by comparing values as number of genes, gene space, etc. to existing annotations for other *Daphnia* genomes. Furthermore, core orthologs from BUSCO’s Arthropoda (odb9) set and conserved domain arrangements from the Arthropoda reference set of DOGMA 3.4 (Dohmen*, et al.* 2016) were searched in the annotated protein set.

The functional annotation was conducted using InterProScan 5.39-77.0 (Jones*, et al.* 2014) as well as a blast against the Swiss-Prot 2019_10.

#### Population samples

##### Raw data QC and contamination check

The quality of raw reads was checked using FastQC v0.11.5. Adapter trimming and quality filtering were performed using Trimmomatic v0.36 with the following parameters: ILLUMINACLIP:TruSeq3-PE.fa:2:30:10 TRAILING:20 SLIDINGWINDOW:4:15 MINLEN:70. For samples sequenced on a NovaSeq6000 instrument and presenting a typical polyG tail, the program fastp (Chen*, et al.* 2018) was used for trimming as well. To assess contamination in the WGA samples FastQ Screen v0.14.0 with the bwa mapping option was used (Wingett and Andrews 2018). A custom database was built to map trimmed reads against possible contaminants that included general common contaminants such as *Homo sapiens*, the UniVec reference database, a bacterial and a viral reference set as well as the *D. galeata* genome and *Acutodesmus obliquus* draft genome (see Supplementary Methods for accession numbers). Samples with <25 % reads mapped to the *D. galeata* genome (and 25% contamination) were excluded from further analysis because the whole amplification of the resting egg most likely failed.

##### Mapping to reference genome and variant calling

The variant calling was performed within the Genome Analysis Toolkit (GATK v4.1.4.0; McKenna*, et al.* 2010) program according to GATK4 best practices (Van der Auwera*, et al.* 2013). The trimmed reads were mapped to the *D. galeata* genome using the BWA-MEM algorithm in BWA v0.7.17 with the -M parameter and adding read group identifiers for Picard compatibility (Li and Durbin 2009). PCR duplicates were marked and filtered out in the BAM file using Picard v2.21.1 (http://broadinstitute.github.io/picard/).

To call variants for each sample GATK HaplotypeCaller in GATK was used with the --emitRefConfidence GVCF option resulting in a genomic variant call format (gVCF) file with information on each position for each individual (Poplin*, et al.* 2018). All gVCF files were consolidated using CombineGVCFs and joint genotyping was performed with GenotypeGVCFs. The VCF file was filtered to include only SNPs and hard filtering was performed to remove variants with a QualByDepth <10, StrandOddsRatio >3, FisherStrand >60, mapping quality <40, MappingQualityRankSumTest <−8 and ReadPosRankSumTest <−5.

Subsequently, we removed sites with either very high coverage (>450) or for which genotypes were missing for more than 20% of the individuals using VCFtools v0.1.5 (Danecek*, et al.* 2011). The final SNP data set for downstream analyses included 3,240,339 SNPs across the 49 samples.

In addition, GenotypeGVCFs was run with the --include-non-variant-sites option to output all variant as well as invariant genotyped sites. The final invariant data set included 127,530,229 sites after removing indels and multi-allelic sites with BCFtools v1.9 (Li 2011) and is used for population genomic analysis to be able to calculate the total number of genotyped sites (variant and invariant) within a genomic window.

As we mapped all different species to the reference *D. galeata* genome we assessed possible reference bias by checking the distribution of reference and alternative alleles observed at heterozygous genotypes based on Pinsky*, et al.* (2021). We pooled all genotypes which were unambiguously assigned to either of the parental species clusters *D. galeata*, *D. cucullata* and *D. longispina* as was done for the population genomic parameters (Table S2) or classified as hybrids using ADMIXTURE inference. Without reference bias, we would expect that in heterozygous genotypes the reference and the alternative allele are on average represented by 50% of the reads. An indication of reference bias would be that the *D. galeata* reference allele would be more frequent.

#### Phylogenetic and population genetics inferences

##### Mitochondrial genome assemblies and phylogenetic analyses

All reads were used to produce mitochondrial genome assemblies using the “de novo assembly” and “find mitochondrial scaffold” modules provided in MitoZ v2.4 with default settings (Meng*, et al.* 2019). For some samples, this was not sufficient and we used two approaches to recover a complete mitogenome: either the mitochondrial baiting and iterative mapping implemented in MITObim v1.9.1 (Hahn*, et al.* 2013) with the *D. galeata* mitochondrial reference genome or the modified baiting and iterative mapping in GetOrganelle v1.7.1 (Jin*, et al.* 2020) with the animal database and k-mer values set to 21, 45, 65, 85 and 105. The procedure used for each dataset is given in Table S5.

We annotated the mitochondrial genome assemblies with the mitochondrial annotation web server MITOS2 (Bernt*, et al.* 2013) using the mitochondrial codon code 05 for invertebrates. Automated genome annotation identified thirteen protein-coding genes (PCGs), two ribosomal RNA genes (rRNAs), and twenty-two transfer RNA genes (tRNAs). Initially, the mitochondrial genes (PCGs and rRNAs, Table S6) were individually aligned with MUSCLE v3.8.1551 (Edgar 2004) and visually checked for their quality. The mitochondrial genome assemblies with discrepancies, i.e., a lot of missing data and/or split features were excluded from further analysis. The final data set (Table S6) included 44 mitochondrial genomes from this study and the previously published mitochondrial genome of *Daphnia laevis* (Martins Ribeiro*, et al.* 2019/, accession number: NC_045243.1/, accession number: NC_045243.1). The mitochondrial genes of the final data set were individually realigned with MUSCLE v3.8.1551 (Edgar 2004) and MACSE v2.05 (Ranwez*, et al.* 2018) and concatenated into a mitochondrial DNA matrix (Table S6) using SequenceMatrix v1.8.1 (Vaidya*, et al.* 2011). During this step, we used MACSE v2.05 to realign PCG genes keeping the information about codon position (gene partitioning) and to remove STOP codons. The final dataset consisted of the concatenation matrix of the thirteen protein-coding (PCGs) and the two structural ribosomal RNA (rRNAs) genes. With this alignment, phylogenetic trees were reconstructed using IQ-TREE v1.6.12 (Nguyen*, et al.* 2015). We initially partitioned the alignment into a full partition model, i.e., each gene and all three codon positions for PCGs, and then ran IQ-TREE with partition analyses (-spp, Chernomor*, et al.* 2016), ModelFinder (-m MFP+MERGE, Kalyaanamoorthy*, et al.* 2017) and 10,000 ultrafast bootstrap (-bb 10000, Hoang*, et al.* 2018) and SH-like approximate likelihood ratio test (-alrt 10000, Guindon*, et al.* 2010) replicates. The resulting trees were visualized in R (R Core Team 2017) using the multifunctional phylogenetics package phytools (Revell 2012).

##### Ancestry and population structure

A principal component analysis was conducted in R v3.6.2 (R Core Team 2017) with the package SNPRelate v1.20.1 (Zheng*, et al.* 2012). Linkage disequilibrium (LD) was calculated within a 500-kb sliding window and LD-pruned for r^2^ values >0.5 before conducting the PCA for all sites using the snpgdsPCA function with default settings. The relative large LD value was chosen because clonal reproduction and the overlap of generations due to diapause leads to increased linkage disequilibrium in *Daphnia* (Brede*, et al.* 2009).

Genetic admixture was estimated using ADMIXTURE v1.3.0 (Alexander and Lange 2011). The SNP set VCF file was converted to BED format using plink v1.90b6.13 (Chang*, et al.* 2015). The log-likelihood values were estimated for one to five genetic clusters (K) of ancestral populations and admixture analysis were run for the most appropriate K value with 10-fold cross-validation. We also conducted the PCA and Admixture analysis using PCAngsd implemented in ANGSD and NgsAdmix, respectively (Korneliussen*, et al.* 2014) to take genotype likelihoods into account (details in Supplementary methods). The results did not differ substantially and are shown in Figure S3.

However, using such a population genetic clustering approach to estimate ancestry coefficients is not directly equivalent to the proportion of hybrid ancestry in each individual and should be interpreted with caution (Kong and Kubatko 2021; Lawson*, et al.* 2018). The results of the ADMIXTURE analysis suggested that the dataset included hybrids between *D. longispina* and *D. galeata* as well as *D. cucullata* and *D. galeata*. We then followed the “ancestry painting” procedure outlined in Barth*, et al.* (2020) and Runemark*, et al.* (2018b), and classified sites according to their FST values when comparing parental species sets. Unlike the PCA and the admixture analysis, this approach requires the user to define parental genotypes; the individuals belonging to these sets are indicated with stars in Figure 1C. Fixed sites are those where a specific allele is fixed in all individuals belonging to one parental species and another allele fixed in the other parental species. To show the ancestry of the hybrid individuals each fixed site was plotted in an “ancestry painting” if at least 80% of genotypes were complete using available ruby scripts (https://github.com/mmatschiner/tutorials/tree/master/analysis_of_introgression_with_snp_data). These scripts calculate the heterozygosity of each individual and visualize regions that are possibly affected by introgression. The mitochondrial genome assembly from each individual was used to determine the maternal species and the proportion of the genome derived from the maternal species was then calculated for each hybrid. For gal x cuc hybrids the hybrid index scale ranges from 0 (gal) to 1 (cuc) and for gal x long hybrids from 0 (long) to 1 (gal).

##### Window-based population parameters

To assess genome-wide genetic differentiation between the clusters identified with admixture, we calculated nucleotide diversity (π), between-taxon differentiation (F_ST_), and between-taxon divergence (d_xy_) using the Python script popgenWindows.py (github.com/simonhmartin/genomics_general release 0.3, Martin*, et al.* 2020) with a sliding 100-kb window, a step size of 10kb and at least 20kb genotyped sites within each window. To compare species pairs we only considered individuals assigned to parental species based on ADMIXTURE results (Table S2). In addition, we also calculated these parameters using one randomly chosen individual from each population per species to check if the estimates are biased because of the overrepresentation of some populations in a species group (Table S2).

Sets of outlier windows were defined as those with F_ST_ values in the upper 95^th^ percentile of the distribution for each of the 3 pairwise comparisons. Further, the genes in these windows were extracted using the annotation file. We used a randomization approach to assess whether the observed intersections (i.e. outlier FST windows occurring in both species) between all seven possible species comparisons are larger or smaller than expected by chance. For this, we randomly drew the observed number of windows, respectively genes from the total number of 10kb windows in the assembly (13,330), respectively the total number of annotated genes (15,845) without replacement and calculated the intersections for all possible comparisons. We compared the resulting intersections from 1,000 replicates with the observed values (Figure S5).

##### Inferring introgression

To identify admixture among three populations we calculated the *f*_3_ statistic with ADMIXTOOLS v 7.0 (Patterson*, et al.* 2012) implemented in the admixr package in R (Petr*, et al.* 2019). We used two parental source populations (A and B) and the target population (C) in the form (C; A, B). Significantly negative *f*_3_ statistic indicates that population C is a mixture of populations A and B or closely related populations.

##### Local ancestry inference

To prepare the SNP set, Beagle v4.1 was used to phase and impute genotypes with 10,000 bp step size and 1000 bp overlapping sliding windows (Browning and Browning 2009). Local ancestry inference was conducted with Loter (Dias-Alves*, et al.* 2018) which infers the origin of each SNP in an admixed individual from two ancestral source populations and doesn’t require additional biological parameters. The respective two parental species populations were used to reconstruct the ancestry tracts of the three putative *galeata* x *cucullata* hybrid individuals and eleven putative *galeata* x *longispina* hybrid individuals using Loter with default settings.

## Authors contributions

MC, JN, AT, SD and MHM performed the sampling and wet lab work. TS, JN and AT conducted the genome assembly and annotation. JN conducted the population genomic analysis and mitochondrial genome reconstruction. TH and MHM conducted the phylogenetic analyses. JN, TS, MC, MP, MHM and TH wrote the manuscript, and all authors edited and contributed to the final version. All authors gave final approval for preprint deposition and publication.

## Data Availability

Genome assembly, annotation and read data (Illumina and PacBio) for the genotype M5 are stored under accession number PRJEB42807. Short read data from resequencing are available in the European Nucleotide Archive under accession numbers ERS5080327-ERS5080375, ERS4993274 and ERS4993282. The annotation used in the present analysis is deposited in Zenodo (DOI: 10.5281/zenodo.4479324), together with supplementary information on the mitochondrial tree (alignment file).

## Acknowledgments

We thank LOEWE-TBG for providing sequencing funds. TH and MM were supported by the grant “SeeWandel: Life in Lake Constance - the past, present and future” within the framework of the Interreg V programme “Alpenrhein-Bodensee-Hochrhein (Germany/Austria/Switzerland/Liechtenstein)”, which funds are provided by the European Regional Development Fund as well as the Swiss Confederation and cantons. MHM was supported by the Austrian Science Fund (FWF): P29667-B25 and J 3774. The funders had no role in study design, data collection and analysis, decision to publish, or preparation of the manuscript. The computational results presented here have been achieved in (part) using the LEO HPC infrastructure of the University of Innsbruck. Some of the data produced and analyzed in this paper were generated in collaboration with the Genetic Diversity Centre (GDC), ETH Zurich.

We thank Jae-Seong Lee and Zhiqiang Ye for giving us access to genome annotations for *Daphnia magna* (SK strain) and *Daphnia pulex* (PA42 strain), respectively. Miklós Bálint provided sediment samples and isotope data for the Arendsee lake. We thank Michael Matschiner for his help with ancestry painting and two anonymous reviewers for their comments on a previous version of the manuscript.

