## Supplementary Methods for "Hybridization dynamics and extensive introgression in the *Daphnia longispina* species complex: new insights from a high-quality *Daphnia galeata* reference genome"

**Supplementary Materials and Methods**

In the following section default options are applied, if not stated otherwise.

**1. Genome assembly**

**Adapter and quality trimming**

Adapter and quality trimming of Illumina reads was conducted with autotrim.pl 0.6.1 (Waldvogel*, et al.* 2018), in combination with Trimmomatic 0.38 (Bolger*, et al.* 2014), FastQC 0.11.7 (Andrews 2010) and MultiQC 1.6 (Ewels*, et al.* 2016) using a custom adapter file and the following Trimmomatic parameters: ILLUMINACLIP: adapter_combined.fa:2:30:10 TRAILING:15 SLIDINGWINDOW:4:20 MINLEN:50.

The tool autotrim.pl allows to trim additionally to adapter and quality for overrepresented *k*-mers which probably arise from smaller parts of adapter sequences.

PacBio subreads were used as provided from the sequencing facility.

**Contamination screening on read level**

To filter out reads possibly originating from contamination, a FastQ Screen like (FQS-like) approach was chosen. In brief, the reads are separated according to mapping behavior to different genomes.

First, a database containing the genomes of *Daphnia magna* (Lee et al., 2019) and *D. pulex* (Ye*, et al.* 2017) as positive controls and the human genome, the genome of the algae the sequenced individuals were fed on as well as several bacterial and viral genomes as negative controls was created. The database contains 108,163 sequences with a total length of 42.4Gb (see Table 1). The accession numbers of the bacterial and viral genomes can be found in the corresponding provided lists.

Table 1: Database parts and corresponding sizes.

| **Species/group** | **Number of sequences** | **Total length [bp]** |
| --- | --- | --- |
| *D. magna* (dmagna-v2.4) | 40,356 | 131,266,987 |
| *D. pulex* (Daphnia_pulex_PA42_v3.0) | 1,822 | 156,418,198 |
| Bacteria | 18,448 | 37,434,584,693 |
| Human (GRCh38.p12) | 594 | 3,257,319,537 |
| *Acutodesmus* *obliquus* (GCA_002149895.1) | 2,707 | 208,176,092 |
| Virus | 44,236 | 1,170,153,228 |

Illumina reads were mapped unpaired (forward and reverse reads separately) with NextGenMap 0.5.5 (Sedlazeck*, et al.* 2013) and the options “‑‑bam 1 ‑‑bin_size 4 ‑‑topn 1000”. PacBio subreads were mapped with minimap2 2.17-r941 (Li 2018) and the options “-H -x map-pb”.

Custom scripts were used to filter reads and to display the results (<https://github.com/schellt/fqs-tools>). The mapping results to the different database parts are displayed in Figure 1. Reads did only pass the filtering if they either did not map to the database at all or had at least one hit against one of the two *Daphnia* genomes. Paired Illumina reads were only returned if both reads pass the filtering. If only one read of a pair passed the filtering they were returned as unpaired. An overview of the effect of different read filtering steps on data volume is shown in Table 2.

**Figure 1:** Mapping results

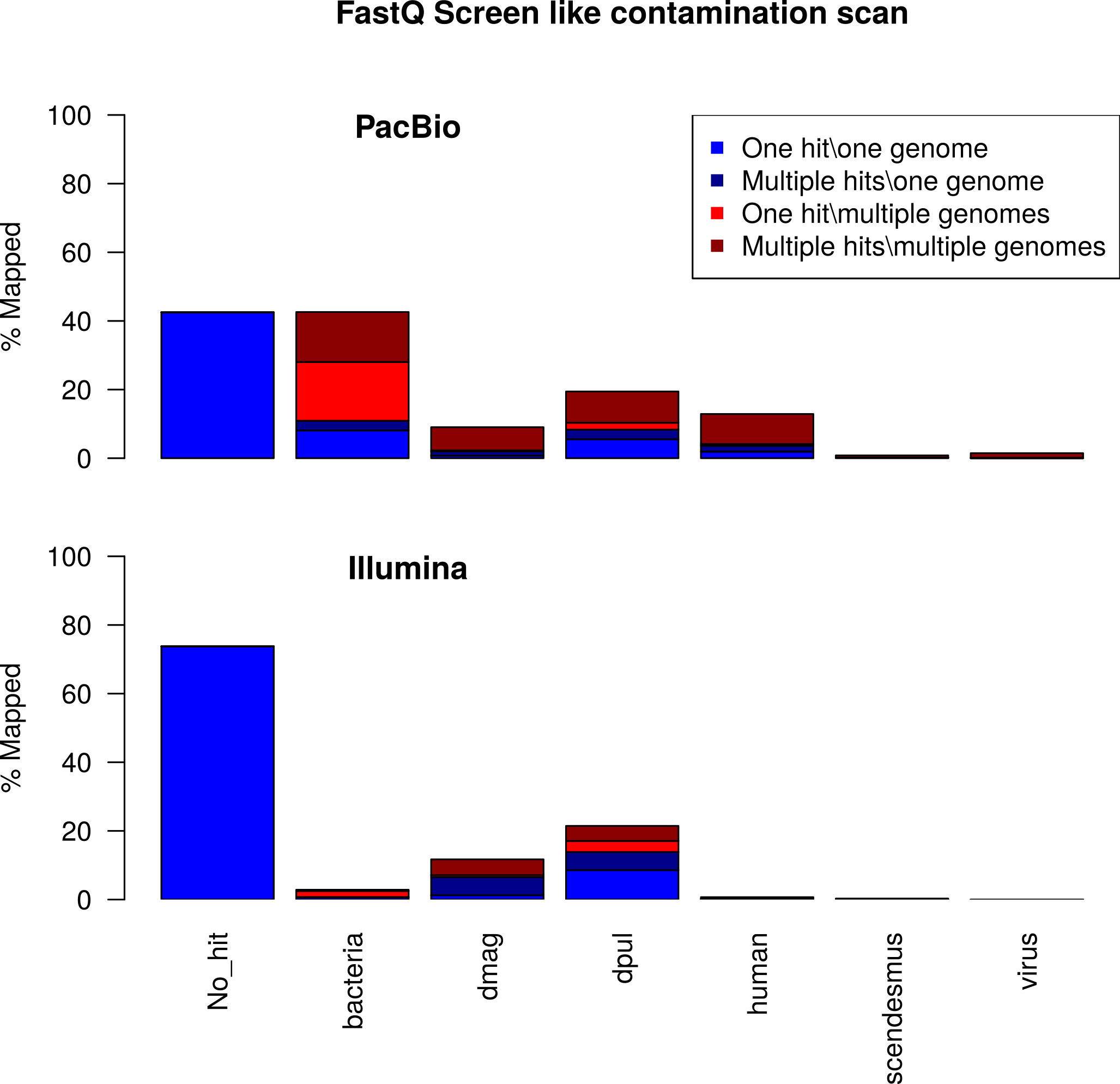

**Table 2:** Comparison of data amount regarding the different preprocessing steps.

|  | | **Raw** | | **Trimmed** | | **FQS-like filtered** | |
| --- | --- | --- | --- | --- | --- | --- | --- |
|  |  | #reads | Gb | #reads | Gb | #reads | Gb |
| Illumina | paired | 77,594,732 | 11.64 | 64,988,122 | 9.13 | 62,533,940 | 8.79 |
|  | unpaired | – | – | 5,322,216 | 0.66 | 5,962,964 | 0.76 |
| PacBio | | – | – | 1,679,290 | 11.52 | 1,121,227 | 6.84 |

**RA assembly**

All paired and unpaired contamination filtered Illumina reads as well as the contamination filtered PacBio reads were used as input for RA 0.2.1 (https://github.com/rvaser/ra).

**Contamination screening on assembly level**

To screen the resulting assembly blobtools 1.0 (Laetsch and Blaxter 2017) was used. The Illumina reads used as input for RA were mapped against the contigs using backmap.pl 0.1 (<https://github.com/schellt/backmap>), which integrates bwa mem 0.7.17-r1188 (Li 2013), samtools 1.9-33-g2d34e15 (Li*, et al.* 2009), Qualimap 2.2.1 (Okonechnikov*, et al.* 2016), bedtools 2.26.0 (Quinlan and Hall 2010) and R 3.5.0 (R Core Team, 2019). To assign Taxonomy IDs blastn 2.9.0+ (Camacho*, et al.* 2009) was used to align the contigs against the complete nt database (-task megablast -outfmt '6 qseqid staxids bitscore' -max_target_seqs 1 -max_hsps 1 -evalue 1e-25). For plotting blobtools 1.1.1 was used.

Contamination with different bacteria can be clearly seen in Figure 2. In total 267 contigs with coverage below 10x and/or GC content above 50% were removed. The removed contigs add up to a total length of 22.97Mb (see detailed Distribution in Figure 3).

To minimize false scaffolding later, PacBio reads mapping to the 267 contigs identified as contamination were removed. This resulted in 871,867 reads with a total length of 5.41Gb which corresponds to 78% and 79% of the FQS-like filtered reads respectively.

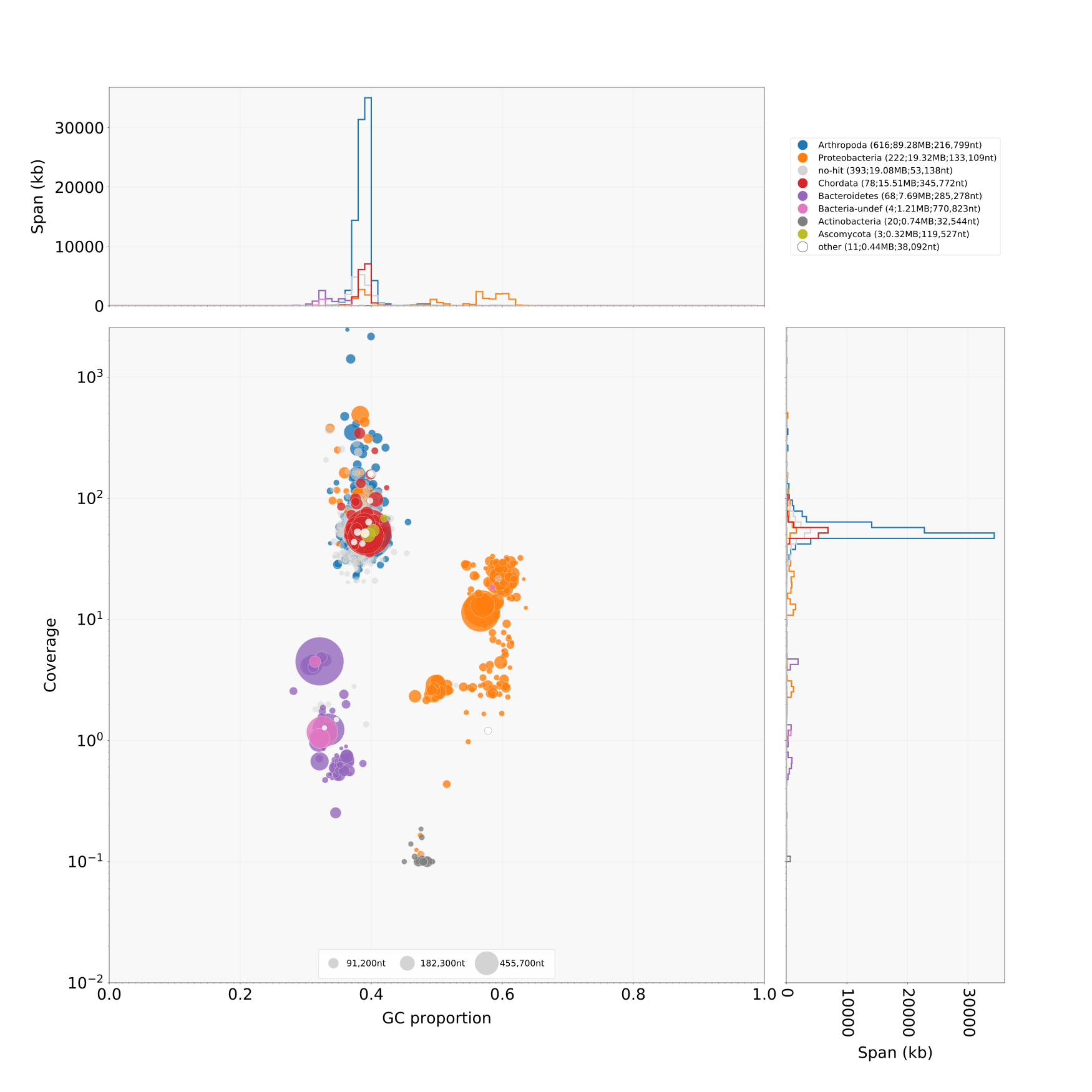

Figure 2: Blobplot of the inital RA assembly. Coverage is based on Illumina data only. Taxonomic assignment was conducted with blastn against the nt database.

**Identifying and removing the mitochondrial genome**

After assembly, the mitochondrial (mt) genome was searched with blastn 2.9.0+ by using the available *D. galeata* mt genomes (accession numbers LC177072.1, LC152879.1, LC177110.1, NC_034297.1, LC177071.1, LC177070.1) as queries against the contigs remaining after blobtools filtering. A total of 15 contigs had hits to at least one of the*se* mt genomes. However, one single contig could be identified by calculating the ratio between alignment length and target sequence (contig) length. The mt genome could be distinguished by the maximum ratio 0.98 from random hits which reached maximally a ratio of 0.008. Finally, one single contig with a length of 15,636bp was removed.

**Scaffolding and gap closing**

Scaffolding and gap closing was conducted in three iterations. Each iteration contains I) a scaffolding with SSPACE LongRead 1-1 (Boetzer and Pirovano 2014), II) gap closing with LR_Gapcloser (<https://github.com/CAFS-bioinformatics/LR_Gapcloser>; commit 156381a) and III) polishing of the former gap parts (that were filled with PacBio reads only) with short reads using wtdbg2-racon-pilon.pl 0.4 (<https://github.com/schellt/wtdbg2-racon-pilon>), which used bwa mem 0.7.17-r1188, samtools 1.9, java 1.8.0_221 (Arnold*, et al.* 2000) and Pilon 1.23 (Walker*, et al.* 2014).

The blobtools filtered PacBio reads were used for scaffolding and gapclosing. Coordinates of gaps produced in the scaffolding step were identified and written into bed format with the help of bedtools 2.28.0. Afterwards the FQS-like filtered paired and unpaired Illumina reads were mapped via bwa mem (-a -c 10000 -t 60) to the gap closed scaffolds. Only reads aligning at least partially to former gap regions were saved to bam format as well as sorted and indexed with samtools. Finally, Pilon was run with additional options “--‑diploid ‑--‑threads 60” on the produced bam files. After running Pilon the scaffold IDs were renamed to match the ones in the bed file again. Short read mapping and Pilon polishing was executed three times in iterative fashion using in all three polishing iterations the gap regions produced at the beginning from the long read‑ scaffolding iteration.

After the third iteration of scaffolding, gap closing and polishing the final scaffolds are yielded.

**Assembly quality assessment**

Contiguity was analyzed with Quast 5.0.2 (Gurevich*, et al.* 2013) at different stages of the assembly process and its main results represented in Table 3.

Table 3: Contiguity statistics of the single assembly steps.

|  | **ra** | **ra-blobfilter-rmmt** | **ra-blobfilter-scaff1** | **ra-blobfilter-scaff2** | **ra-blobfilter-scaff3** |
| --- | --- | --- | --- | --- | --- |
| #Sequences | 1,415 | 1,147 | 473 | 370 | 346 |
| Total length [Mb] | 153.6 | 130.6 | 132.9 | 133.2 | 133.3 |
| N50 [kb] | 172.4 | 175.3 | 533.0 | 729.4 | 756.7 |
| GC [%] | 40.02 | 38.74 | 38.75 | 38.75 | 38.75 |
| N’s [%] | ~0 | ~0 | 0.07 | 0.09 | 0.09 |

To look at mapping rate, coverage and insert size distribution, etc., backmap 0.3 (<https://github.com/schellt/backmap>) was used in combination with bwa mem 0.7.17-r1188, Minimap 2.17-r941, Samtools 1.9, Qualimap 2.2.1, bedtools 2.28.0, R 3.5.3 and MultiQC 1.8. Mapping the paired and unpaired FQS-like filtered Illumina reads and the blobtools filtered PacBio reads resulted in a mapping rate of 94.1% and 85.5%, respectively, according to Qualimap’s bamqc. The insert size of the paired Illumina reads is narrowly distributed around a median of 327 (Figure 3). The genome size was estimated based on mapped nucleotides and mode of the coverage distribution by backmap, resulting in 156.86Mb and 178.03Mb for Illumina (52x) and PacBio (26x) respectively. Additionally, the genome size estimated using a k-mer based approach with GenomeScope (150.6Mb) can be found here: <http://qb.cshl.edu/genomescope/analysis.php?code=WeYH4KGn4W7eTjIa1qhf>.

To show absence of contamination in the assembly a blastn search of the final scaffolds against the nt database was conducted as above, and the results plotted with blobtools 1.1.1.

Completeness in terms of single copy core orthologs of the final scaffolds was assessed with BUSCO 3.0.2 (Simão*, et al.* 2015), using the Arthropoda set (odb9) and the option --‑long. This resulted in C: 95.7% [S: 94.7%, D: 1.0%], F: 0.8%, M: 3.5%, n‑: 1066.

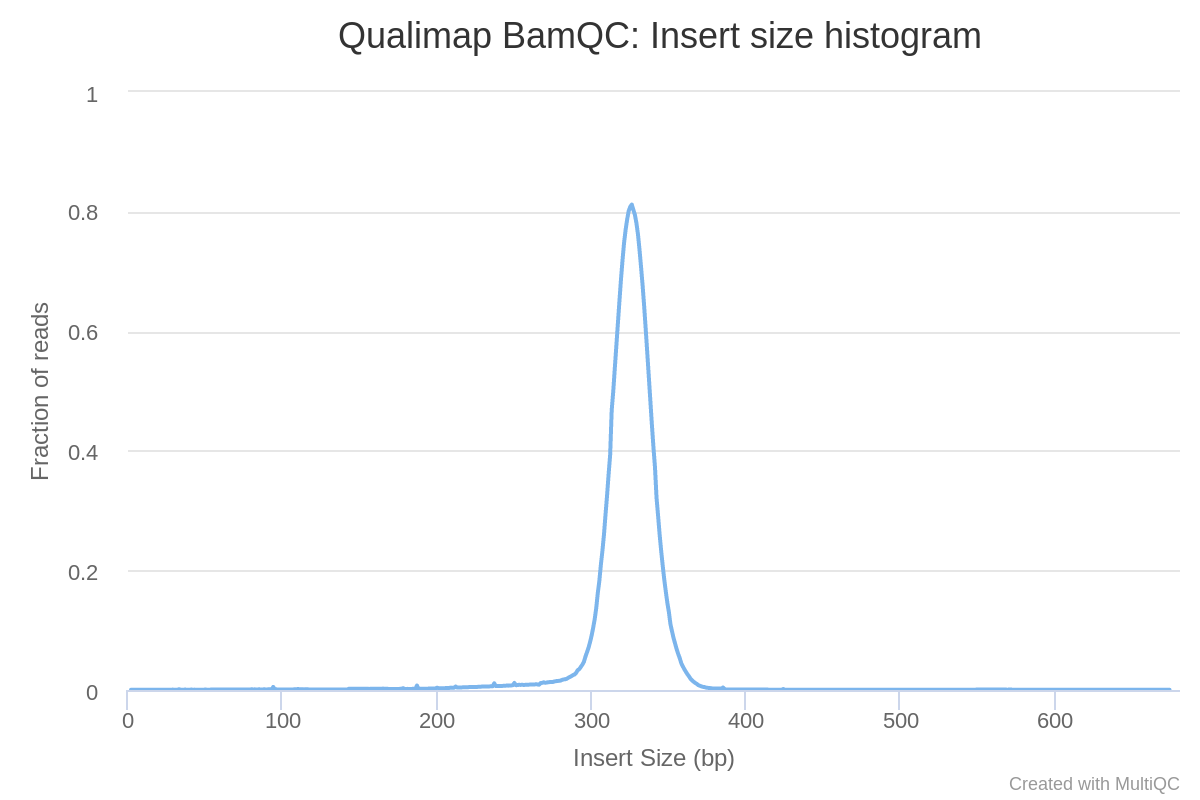

Figure 3: Insert size distribution of paired Illumina reads. Created with Qualimap and MultiQC.

**2. Genome annotation**

Before starting with the annotation the fasta headers of the genome assembly and the *D. galeata* transcriptome assembly (HAFN01.1) were simplified with Augustus’ simplifyFastaHeaders.pl.

**Repeat library creation and repeat masking**

To identify *D. galeata* specific repeats, RepeatModeler 2.0 (Smit and Hubley 2015) in combination with RepeatMasker 4.1.0 (Smit*, et al.* 2013-2015) including RepBase release 20181026 (Bao*, et al.* 2015), RECON 1.08 (Bao and Eddy 2002), RepeatScout 1.0.6 (Price*, et al.* 2005), Tandem Repeats Finder 4.0.9 (Benson 1999) and RMBlast 2.9.0+. RepeatModeler was run with the options “-pa 10 -LTRStruct” and resulted in 1,115 families with a total length of ~1Mb. Detailed distribution of repeat family classification can be seen in Table 4.

Table 4: Distribution of repeat family classification.

| **Classification** | **Number of families** | **Total length [bp]** |
| --- | --- | --- |
| Unknown | 796 | 474,907 |
| LTR | 145 | 249,242 |
| DNA | 93 | 155,424 |
| LINE | 35 | 70,649 |
| tRNA | 31 | 43,674 |
| SINE | 13 | 17,071 |
| RC | 5 | 7,080 |
| Simple repeat | 4 | 364 |
| SINE | 2 | 5,442 |
| snRNA | 1 | 3,215 |

The yielded repeat families were combined with 237 *D. pulex* and 1 *D. pulicaria* repeat sequences from RepBase release 20181026 to create the final repeat library with 1,353 sequences and a total length of 1.78Mb.

The genome assembly was then soft masked for later usage within Augustus with the final repeat library. To do so, RepeatMasker 4.1.0 was applied with the search engine rmblastn 2.9.0+, Tandem Repeats Finder 4.0.9 and the additional options “-xsmall -no_is -e ncbi -pa 10 -s”. Afterwards 21.9% of the assembly were masked. The distribution of masked fraction per repeat element can be found in Table 5.

Table 5: Repeat masking results as provided from RepeatMasker. Most repeats fragmented by insertions or deletions have been counted as one element.

| **Classification** | **Number of elements** | **Percentage of assembly** |
| --- | --- | --- |
| Unknown | 48,249 | 10.18 |
| LTR | 7,824 | 5.25 |
| DNA | 7,145 | 2.29 |
| Simple repeat | 60,548 | 1.58 |
| LINE | 2,631 | 1.36 |
| snRNA | 1,591 | 0.49 |
| Low complexity | 12,696 | 0.43 |
| SINE | 849 | 0.19 |
| RC | 444 | 0.13 |

**Creation of initial gene prediction models**

Three different gene prediction models were produced with Augustus, GeneMark and SNAP. The detailed procedure is described below.

One model was created using autoAug.pl from Augustus 3.3.2 (Stanke*, et al.* 2008) in combination with PASA 2.4.1 (Haas*, et al.* 2003) and GMAP 2019-09-12 (Wu and Watanabe 2005). As Input for autoAug.pl were used the soft masked assembly and the *D. galeata* transcriptome (HAFN01.1, Huylmans*, et al.* 2016) as well as the additional options “-v ‑-v ‑-v ‑--‑pasa ‑--‑useGMAPforPASA ‑--‑‑noninteractive”.

A second model was created using GeneMark ET 4.48_3.60_lic (Lomsadze*, et al.* 2005). First different RNAseq reads (ERR1551794 and ERR1551390- ERR1551397) were trimmed with autotrim.pl. The trimmed RNAseq reads where mapped paired and unpaired separately with HISAT 2.1.0 (Kim*, et al.* 2019) and the additional option “-p 70”. The two resulting bam files were merged and sorted (‑-l 9 ‑-@ 10) using samtools 1.9. Afterwards Augustus’ bam2hints with the additional options “‑--‑minintronlen=20 ‑--‑maxintronlen=500000” was used to produce a gff file containing possible introns. This gff file was filtered using filterIntronsFindStrand.pl (included in Augustus) with the additional option “‑--‑score”. Finally, gmes_petap.pl from GeneMark was run with the unmasked genome assembly, the filtered intron gff file and the additional options “‑--‑v ‑--‑cores=88 ‑--‑‑max_intron=400000”.

A third model was created with busco2snap.pl 0.1 (<https://github.com/schellt/busco2snap>) in combination with SNAP 2006-07-28 (Korf 2004), Augustus 3.3.2 and blastp 2.9.0+. In brief, the intron-exon-boundaries of 1010 complete and single copy BUSCOs are predicted with the Augustus model created with BUSCO. If more than one gene is predicted per locus a blastp search against the ancestral variants of the BUSCO genes is used to select the most likely one. Afterwards the exon annotation is converted from gff to zff and finally the SNAP model is created with its tools fathom, forge and hmm-assembler.pl.

**Structural annotation**

The structural annotation was conducted in MAKER 2.31.10 (Holt and Yandell 2011) in combination with blast 2.10.0+, RepeatMasker 4.1.0, Exonerate 2.4.0 (Slater and Birney 2005), SNAP 2013-11-29, GeneMark 3.60_lic, Augustus 3.3.2 and tRNAscan-SE 1.3.1 (Lowe and Eddy 1997). The following input sequences were used for MAKER: the unmasked genome assembly, the species own transcriptome assembly (HAFN01.1) as ESTs, the complete Swiss-Prot 2019_10 (Consortium 2019), and the protein sequences resulting from *D. magna* (Lee*, et al.* 2019) as well as *D. pulex* (Ye*, et al.* 2017) genome annotations as protein evidence. Repeat masking was conducted with the above described final repeat library for RepeatMasker and MAKER’s te_proteins.fasta for RepeatRunner. For model based gene prediction the above described models from SNAP, GeneMark and Augustus were fed into MAKER. The options est2genome, protein2genome, trna and alt_splice were switched on. The minimum protein length was set to 10aa. MAKER was compiled and executed with mpich 3.3.2 and the additional option “-fix_nucleotides”. With this, the first round of MAKER was completed.

Afterwards one gff file with and one without the assembly sequence were created using MAKER’s gff3_merge. From the gff without assembly sequence the tracks “est2genome”, “protein2genome” and “repeatmasker”/”repeatrunner” were extracted and saved in three separate gff files.

The gff file with assembly sequence was used as input for MAKER’s maker2zff. The resulting files were processed with fathom (fathom genome.ann genome.dna -categorize 1000 && fathom -export 1000 -plus uni.ann uni.dna), forge (forge export.ann export.dna) and hmm-assembler.pl to create a new SNAP model based on the first round of MAKER.

The Augustus model was retrained with autoAug.pl as above, except that the gff file without assembly sequence from first round’s results was specified as trainingset.

The second round of MAKER was conducted as the first round, but instead of specifying the same evidence and repeat sequences again, the gff files containing the tracks est2genome, protein2genome and repeats as est_gff, protein_gff and rm_gff were used respectively. Furthermore, the retrained/new models from Augustus and SNAP were used.

After finishing the second round of MAKER, the Augustus and SNAP model were retrained/newly created as above.

The third and final round of MAKER was run as the second one, except of updating the Augustus and SNAP model. The options est2genome and protein2genome were switched off, whereas the option always_complete was switched on.

Finally, a gff file containing the maker track only without the assembly sequence was created with gff3_merge. The transcripts and protein fasta files of the annotated genes were created with fasta_merge.

**Structural annotation quality assessment**

The final gff file was processed with a custom script to count and calculate several contiguity statistics of the structural annotation except tRNAscan results. Furthermore, the annotated protein set was analyzed using BUSCO 3.0.2 in combination with the Arthropoda set (odb9) and DOGMA 3.4 (Dohmen*, et al.* 2016) in combination with Pfam scan 1.6 and DOGMA’s reference set for Arthropoda. The results are compared in Table 6.

Table 6: Structural annotation statistics. Contiguity statistics of the annotation were calculated excluding tRNAscan results. BUSCO 3.0.2 was executed in protein mode for the different MAKER rounds. Conserved Domain Arrangements (CDAs) were searched with Pfam scan 1.6 and DOGMA 3.4.

|  | **Assembly** | **Round 1** | **Round 2** | **Round 3** |
| --- | --- | --- | --- | --- |
| Number |  |  |  |  |
| Gene |  | 15513 | 15429 | 15845 |
| mRNA |  | 16471 | 16275 | 16774 |
| Exon |  | 121995 | 119033 | 117364 |
| CDS |  | 130242 | 123265 | 119402 |
| Mean number of |  |  |  |  |
| mRNAs/gene |  | 1.06 | 1.05 | 1.06 |
| Exons/mRNA |  | 7.41 | 7.31 | 7.00 |
| CDSs/mRNA |  | 7.91 | 7.57 | 7.12 |
| Median length |  |  |  |  |
| Gene |  | 2145 | 2187 | 2097 |
| mRNA |  | 2236 | 2248 | 2142 |
| Exon |  | 163 | 164 | 167 |
| Intron |  | 74 | 74 | 74 |
| CDS |  | 150 | 151 | 152 |
| Total space |  |  |  |  |
| Gene |  | 54765381 | 52605643 | 51689473 |
| mRNA |  | 54765288 | 52605588 | 51689329 |
| Exon |  | 29662778 | 29222288 | 29314592 |
| CDS |  | 25750445 | 25271022 | 25132876 |
| Single |  |  |  |  |
| Exon mRNA |  | 1015 | 746 | 663 |
| CDS mRNA |  | 1104 | 854 | 710 |
| BUSCO [%] N=1066 |  |  |  |  |
| Complete | 95,7 | 93,9 | 93,9 | 94,3 |
| Single copy | 94,7 | 91,0 | 91,0 | 91,7 |
| Duplicated | 1,0 | 2,9 | 2,9 | 2,6 |
| Fragmented | 0,8 | 0,8 | 1,0 | 0,7 |
| Missing | 3,5 | 5,3 | 5,1 | 5,0 |
| CDAs [%] N=4222 |  | 93,94 | 93,89 | 93,63 |

As further quality criterion the Annotation Editing Distance (AED) was compared between the three MAKER rounds and is visualized in Figure 4.

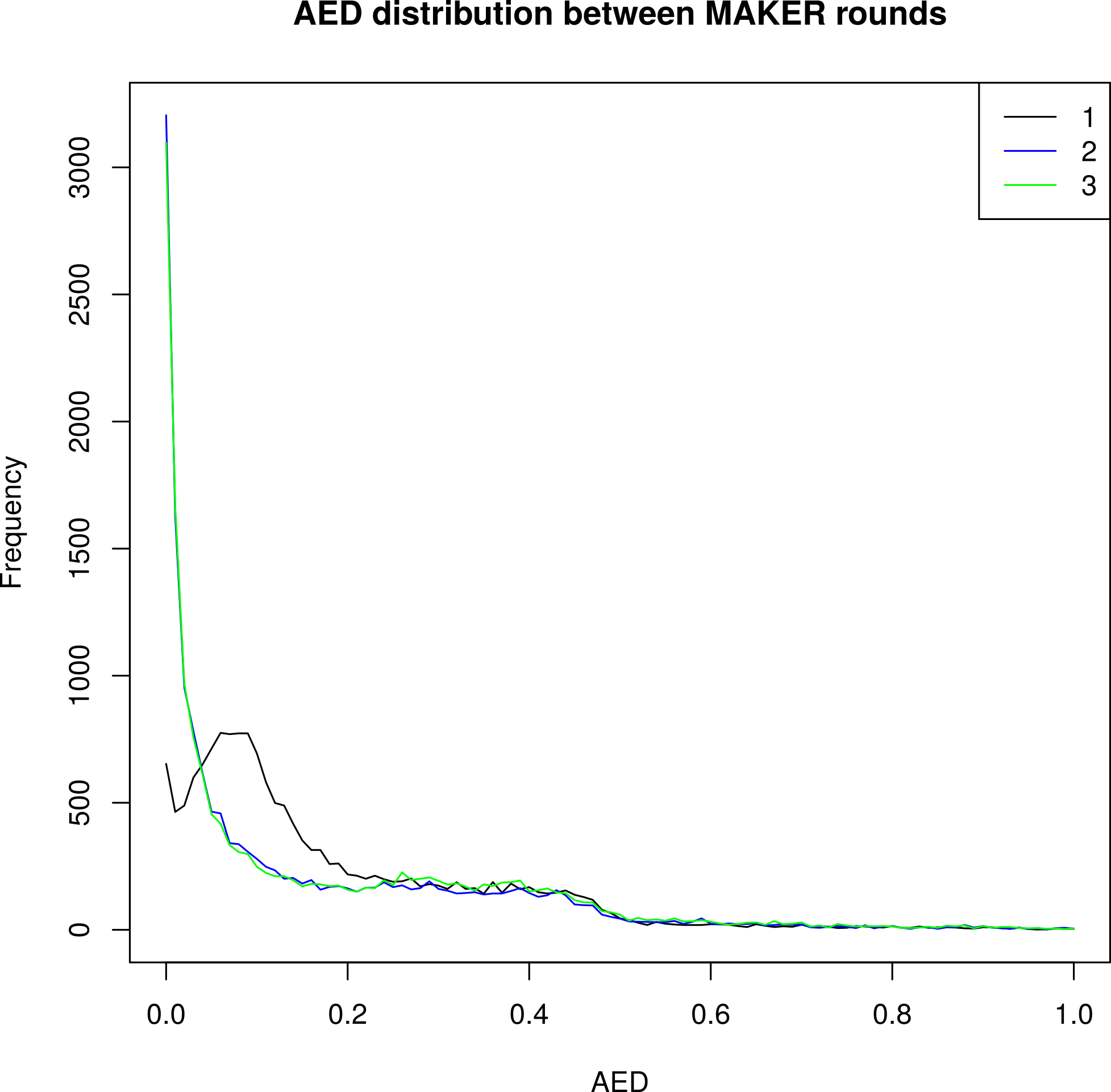

Figure 4: Annotation Editing Distance (AED) distribution of the different MAKER iterations. The AED shows congruency between the annotation and evidence alignments. Low values indicate high congruency.

**Functional annotation**

The functional annotation of the annotated protein set was conducted using InterProScan 5.39-77.0 (Jones*, et al.* 2014) in combination with the databases/analyses CDD 3.17 (Lu*, et al.* 2020), Coils 2.2.1 (Lupas*, et al.* 1991), Gene3D 4.2.0 (Yeats*, et al.* 2006), Hamap 2019_01 (Pedruzzi*, et al.* 2015), MobiDBLite 2.0 (Necci*, et al.* 2017), PANTHER 14.1 (Mi*, et al.* 2012), Pfam 32.0 (El-Gebali*, et al.* 2019), Phobius 1.01 (Käll*, et al.* 2004), PIRSF 3.02 (Wu*, et al.* 2004), PRINTS 42.0 (Attwood*, et al.* 2012), ProSitePatterns 2019_01 and ProSiteProfiles 2019_01 (Sigrist*, et al.* 2012), SFLD 4 (Akiva*, et al.* 2014), SignalP_EUK 4.1 (Nielsen 2017), SMART 7.1 (Letunic and Bork 2018), SUPERFAMILY 1.75 (Wilson*, et al.* 2009), TIGRFAM 15.0 (Haft*, et al.* 2001) and TMHMM 2.0c (Krogh*, et al.* 2001). The lookup for pathways and Gene Ontology (GO) terms was switched on.

To additionally assign putative names, an alignment of the annotated protein set against the Swiss-Prot 2019_10 was conducted using blastp 2.9.0+ with the options “‑num_threads 70 ‑max_hsps 1 ‑max_target_seqs 1 ‑outfmt 6”

.

In total 15,898 (94.78%) and 15,960 (95.15%) protein sequences could be annotated by InterProScan and blast against Swiss-Prot, respectively. In combination, a functional annotation for 16,675 (99.41%) protein sequences could be achieved. A detailed overview of the functional annotated sequences per database or search algorithm is shown in Table 7.

Table 7: Functional annotation success of the different Databases and search algorithms.

| **Database/analysis** | **Number of different protein sequences** |
| --- | --- |
| **Interpro** |  |
| CDD | 5550 |
| Coils | 3191 |
| Gene3D | 9977 |
| Hamap | 279 |
| MobiDBLite | 6661 |
| PANTHER | 13040 |
| Pfam | 11567 |
| Phobius | 6876 |
| PIRSF | 793 |
| PRINTS | 2565 |
| ProSitePatterns | 3169 |
| ProSiteProfiles | 6378 |
| SFLD | 84 |
| SignalP_EUK | 2951 |
| SMART | 5444 |
| SUPERFAMILY | 9816 |
| TIGRFAM | 721 |
| TMHMM | 4211 |
| GO | 9555 |
| Reactome | 3927 |
| **Total** | **15898** |
| **Total [%]** | **94.78** |
| **blast** |  |
| Swiss-Prot | 15960 |
| Swiss-Prot [%] | 95.15 |
| **Overall total** | **16737** |
| **Overall total [%]** | **99.78** |

The results of the functional annotation was then included in the gff and the transcript/protein fasta files using adapted versions of MAKER’s maker_functional_gff and maker_functional_fasta respectively (<https://github.com/schellt/maker-functional>).

**ANGSD**

To confirm the result of the GATK based variant calling, we estimated genotype likelihoods using ANGSD v0.931-2-gfd2a527 with the following parameters: angsd -P 32 -bam $BAM -ref $REF -out $OUT -GL 2 -doGlf 2 -doMajorMinor 1 -doMaf 1 -SNP_pval 1e-6 -uniqueOnly 1 -remove_bads 1 -only_proper_pairs 1 -trim 0 -C 50 -baq 1 -minMapQ 20 -minQ 20 -setMinDepth 400 -setMaxDepth 2000 -minMaf 0.05 -doCounts 1 and performed admixture analysis using NGSadmix and PCA analysis using PCAngsd v1.02.
