## Supplementary Figures for "Hybridization dynamics and extensive introgression in the *Daphnia longispina* species complex: new insights from a high-quality *Daphnia galeata* reference genome"

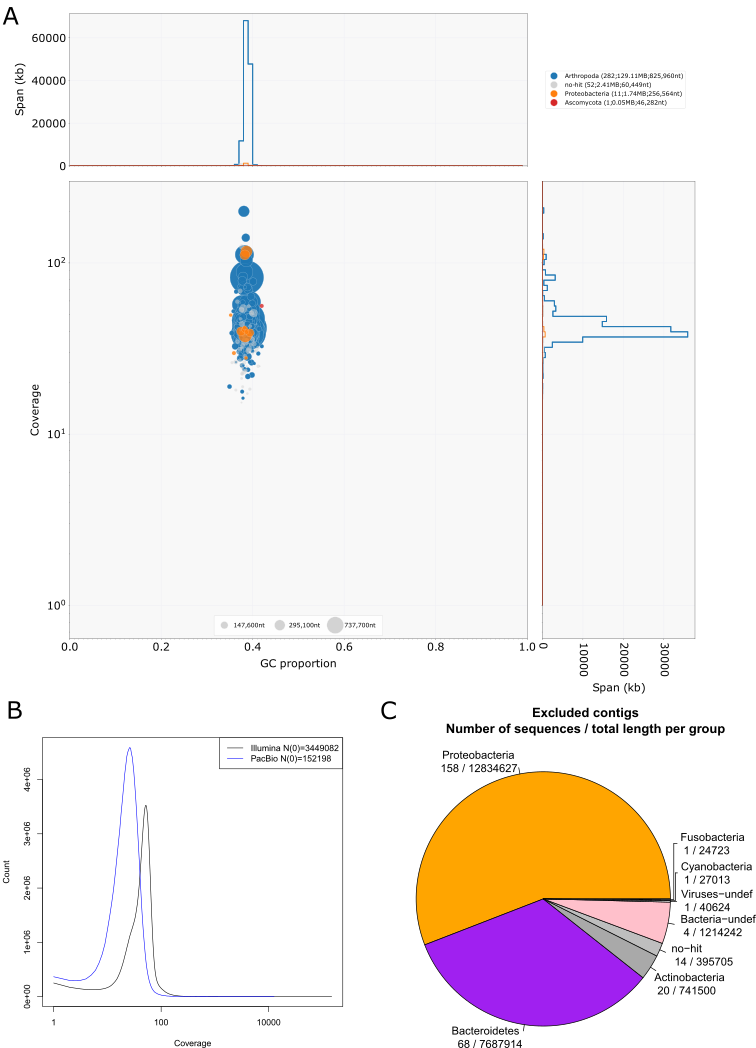

**Figure S1:**  Assembly quality assessment- **A**. Blobplot of the final assembly. Coverage is based on Illumina and PacBio data. Taxonomic assignment was conducted with blastn against the nt database. **B**. Coverage distribution per sequencing technology of the final assembly **C**. Excluded contigs. The pie chart displays the total length of the excluded contigs per taxonomic group. Below the group name the number of contigs and their total length is specified.

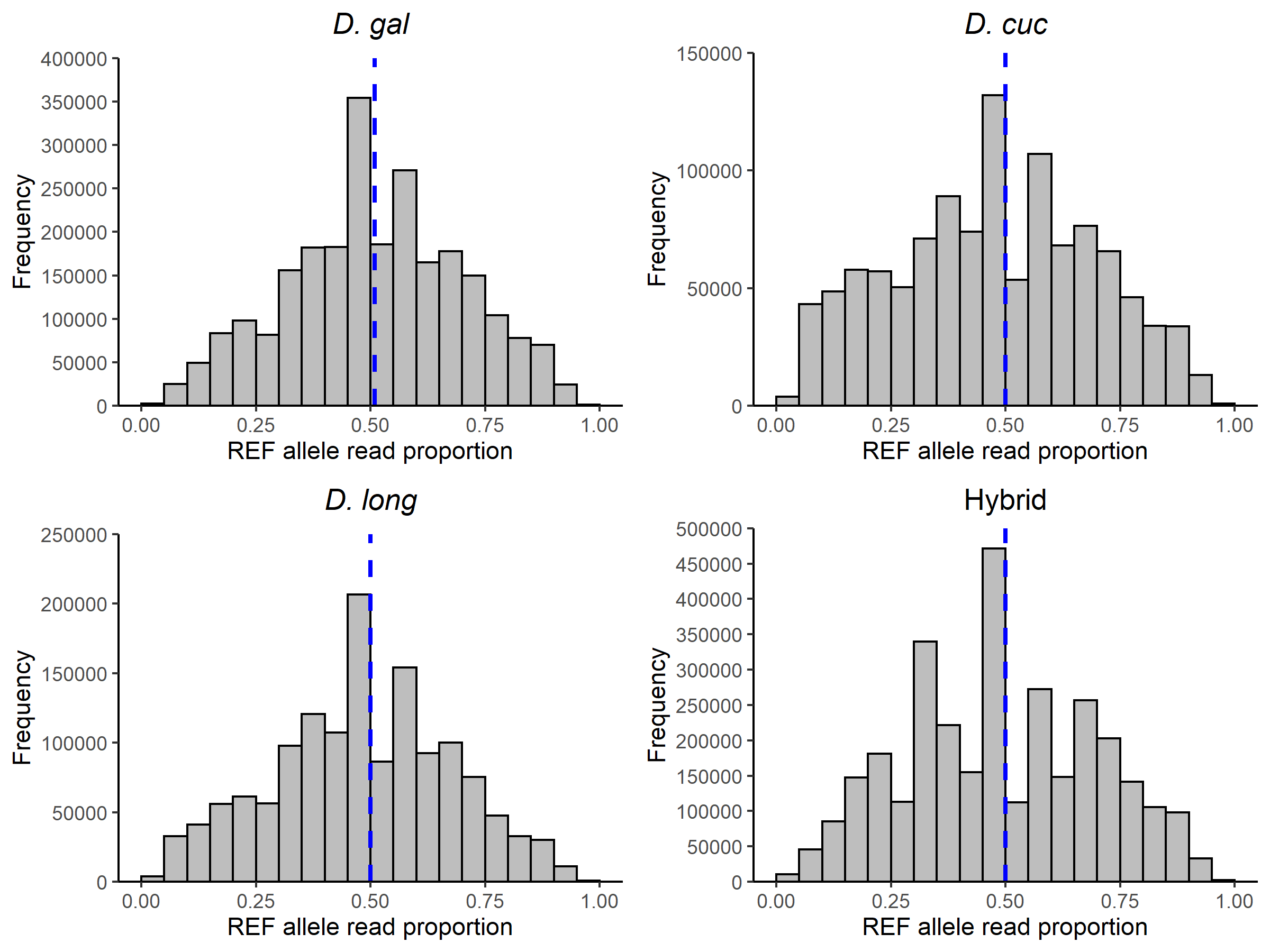

**Figure S2:** Distribution of reference and alternative alleles observed at heterozygous genotypes for all individuals which were unambiguously assigned to either of the parental species clusters *D. galeata*, *D. cucullata* or *D. longispina* or classified as hybrids. The centring of this distribution around the blue median line suggests that there is no biased mapping of the reference alleles.

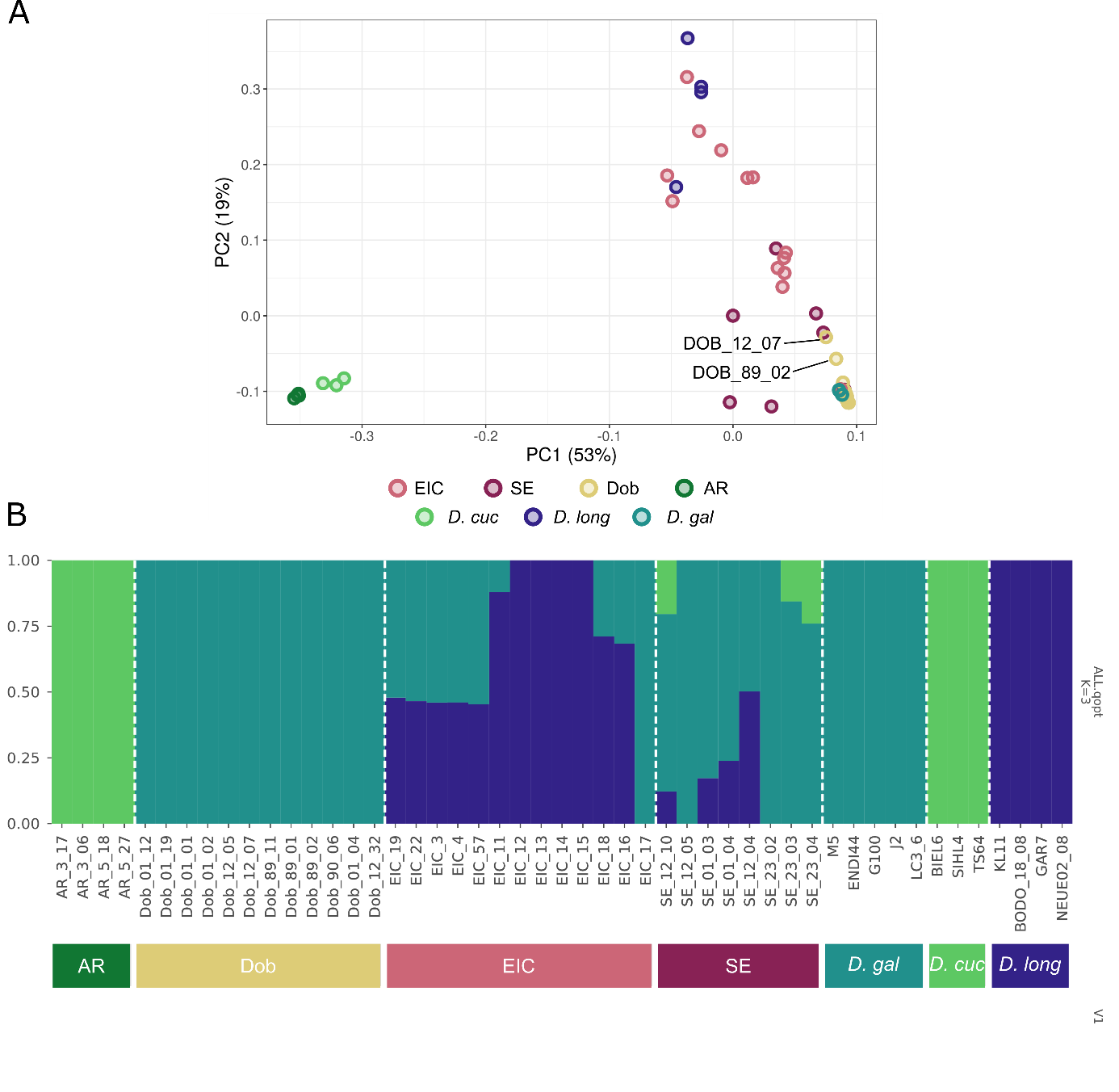

**Figure S3: A.** PCA plot obtained with PCAngsd. **B.** Admixture plot obtained with K=3 in NgsAdmix. *D.gal:* *D. galeata*, *D. long*: *D. longispina*, *D. cuc*; *D. cucullata*. Bottom bars are color coded to match the color scheme used in panel A.

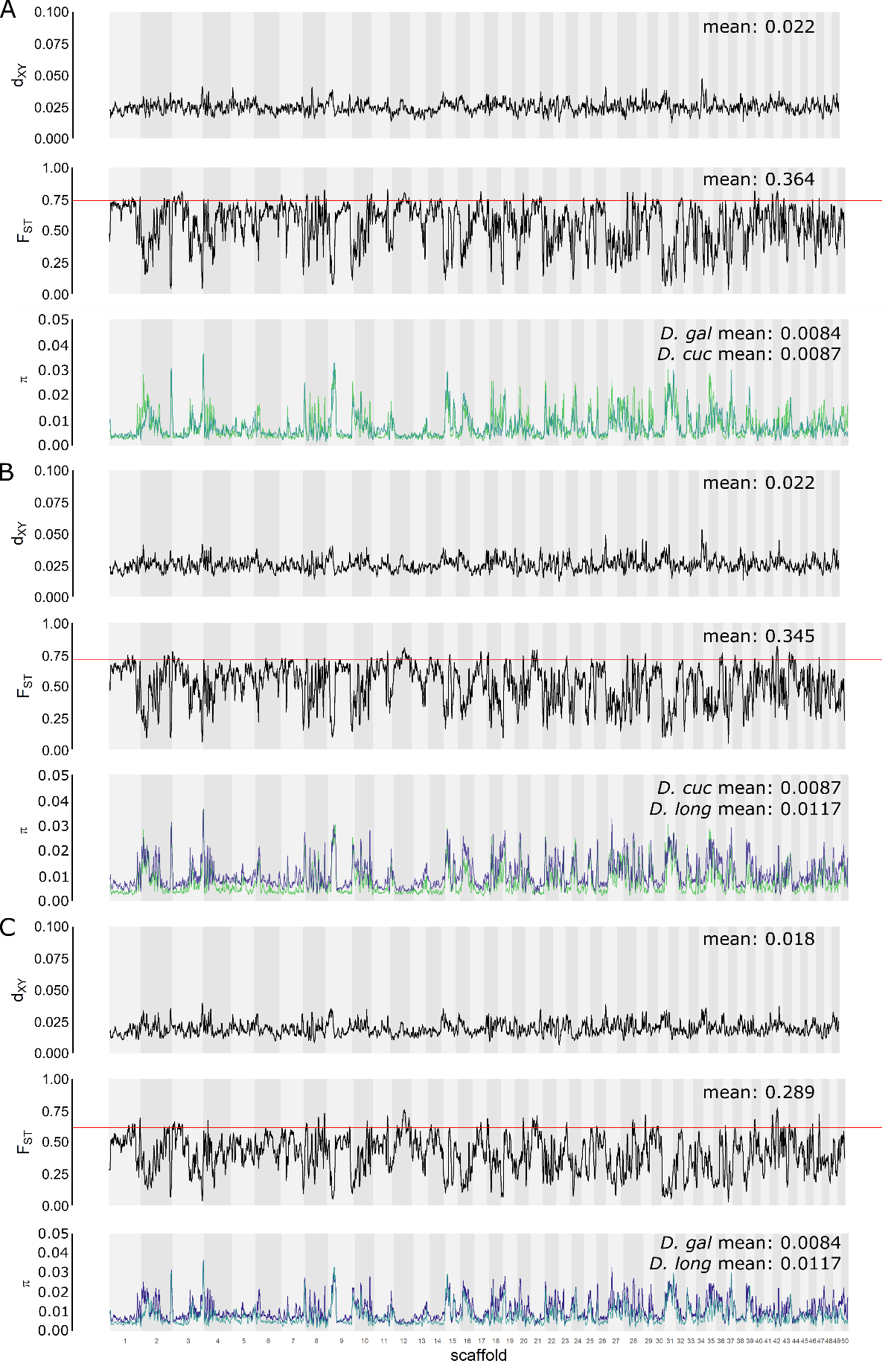

**Figure S4:** Window-based statistics reduced to one individual per population for the pairs **A**. *D. galeata*/ *D. cucullata,* **B**. *D cucullata*/ *D. longispina* and **C**. *D. galeata*/ *D. longispina*, shown for the 50 largest scaffolds in 100kb windows with 10kb step size – calculations are for one randomly chosen individual representing each population per species. **In each panel from top to bottom:** d_xy_ values, pairwise F_ST_ values with a red horizontal line indicating the 95^th^ percentile, nucleotide diversity (π) for *D. galeata* (teal), *D. longispina* (dark blue) and *D. cucullata* (lime green).

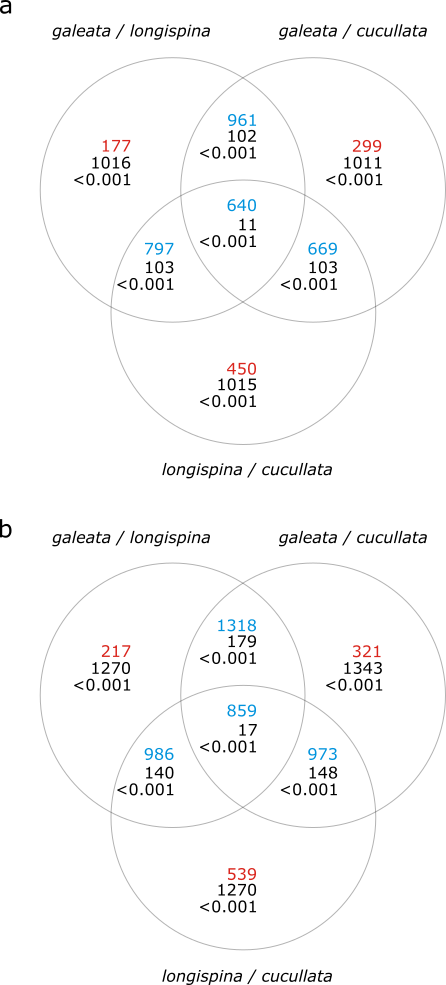

**Figure S5:** Venn-diagram for **a**. outlier windows (based on *F_ST_* values) **b**. genes within these outlier windows. The upper number in each region of the diagram is the observed number of items in the region, the middle number the mean expected number according to 1000 randomisations and the lower number gives the probability of this outcome. The observed number is red when smaller than expected and blue when larger. *galeata / longispina*: set of elements found in the comparison between *D. galeata* and *D. longispina*, Figure 3C. *galeata / cucullata*: set of elements found in the comparison between *D. galeata* and *D. cucullata*, Figure 3A. *longispina / cucullata*: set of elements found in the comparison between *D. longispina* and *D. cucullata*, Figure 3B.

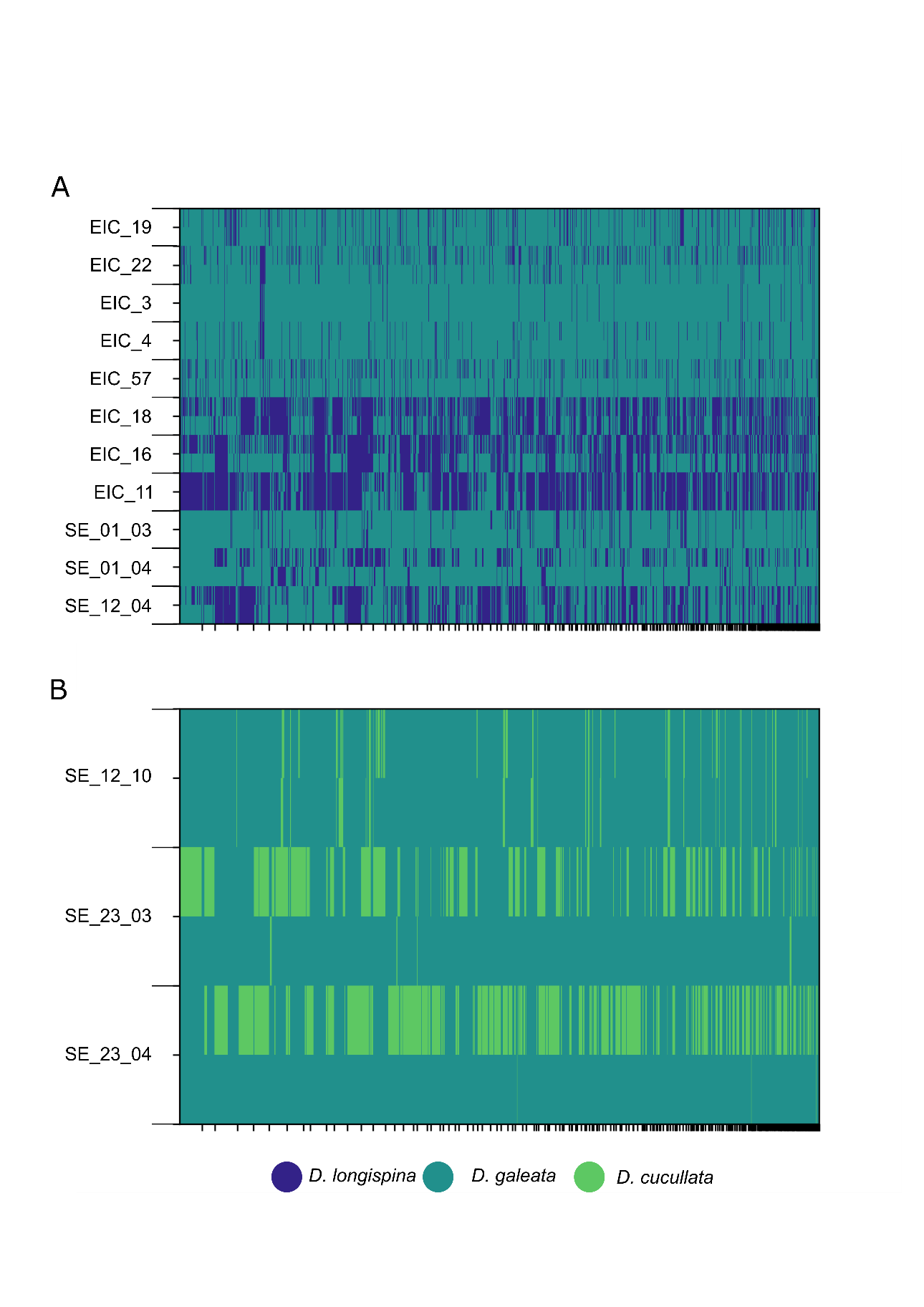

**Figure S6:** Local ancestry inference for: **A.** D. galeata x longispina individuals. **B.** 3 D. galeata x cucullata individuals. Ancestry tracts are visualized on the y-axis in two rows for the two haplotypes of one individual and on the x-axis along the genomic scaffolds.
